# Ecological corridors homogenize plant root endospheric mycobiota

**DOI:** 10.1101/2022.09.02.506380

**Authors:** Jie Hu, Philippe Vandenkoornhuyse, Fadwa Khalfallah, Romain Causse-Védrines, Cendrine Mony

## Abstract

Ecological corridors have been shown to promote species coexistence in fragmented habitats where dispersal limits species fluxes. The corridor concept was developed and investigated mainly by focusing on macroorganisms while microorganisms, the invisible majority of biodiversity, have been disregarded.

Combining an experimental corridor-mesocosm design with high-throughput amplicon sequencing, we analyzed the effect of corridors on the dynamics of endospheric fungal assemblages associated with plant roots at metric scale over two years (i.e. at five time points).

We show that the plant symbiotic compartment was sensitive to corridor effects when the corridors were set up at a small spatial scale. The endospheric mycobiota of connected plants displayed higher species richness, lower beta-diversity, and a more deterministic assembly than the mycobiota of isolated plants. These effects became more pronounced with the development of host plants.

Biotic corridors composed of host plants may thus play a key role in the spatial dynamics of microbial community and may influence microbial diversity and related ecological functions.

## Introduction

Anthropogenic activities have caused habitat destruction and degradation leading to habitat fragmentation and threatening biodiversity (Diamond, 1976; Hilty JA *et al.*, 2006; Haddad *et al.*, 2015). Ecological corridors have been considered as one way to mitigate the negative effect of habitat fragmentation (Rosenberg *et al.*, 1997; Haddad, 2015). When dispersal among habitat patches is limited, corridors facilitate the movements of animals and the dispersal of plants between habitat patches (Tewksbury *et al.*, 2002) and, in what is known as the rescue effect, reduce the probability of local extinction of isolated populations known as the rescue effect (Brown & Kodric-Brown, 1977). Corridors therefore have mostly positive effects on species richness at the patch scale (Damschen, 2006) although there may be some negative effects such as also facilitating the dispersal of highly competitive species or pathogens (Haddad *et al.*, 2014). In metacommunities, which are defined as a set of connected patches linked by dispersal fluxes, local community assembly involves both deterministic and stochastic processes, competition, drift, and dispersal being the key elements (Leibold *et al.*, 2004). The metacommunity framework predicts that species local diversity will increase when the dispersal rate is intermediate (hump-shaped relationship among dispersal effects and local diversity), while patches with high dispersal fluxes are thought to be less subjected to drift, and to harbor more similar communities (Mouquet & Loreau, 2003). Beta-diversity among local communities is therefore reduced (Jamoneau *et al.*, 2012) due to high migration rates (Arévalo *et al.*, 2010). Yet, research on the effect of ecological corridors on maintaining biodiversity has mostly focused on macroorganisms, while microorganisms, the invisible majority of biodiversity, have mainly been overlooked. Existing knowledge on ecological corridors applied to microorganisms focuses on either key pathogen species or on specific taxonomic groups such as mycorrhizal fungi (Groppe *et al.*, 2001; Yuen & Mila, 2015; Chagnon *et al.*, 2020), but does not account for the diversity, composition and assembly process of the microbial assemblage as a whole.

Microbial distribution was long assumed to be subject to very little dispersal limitation due to small size and high propagule production (Baas-Becking’s hypothesis) (Baas Becking, L. G. M., 1934), therefore depending only on local habitat conditions. This assumption was contradicted by evidence for biogeographical patterns in microbes (Martiny *et al.*, 2006; Hanson *et al.*, 2012; Nemergut *et al.*, 2013; Xu *et al.*, 2022) and strong distance-decay relationships (Xu *et al.*, 2021), while mechanistic approaches, based on the metacommunity framework quantified the importance of dispersal in species coexistence in microbes (Miller *et al.*, 2018; Langenheder & Lindström, 2019). The more recent microbial community coalescence concept (Rillig et al., 2015) suggests that entire local microbial communities may disperse among habitats either via dispersal vectors or habitat mixing. Most studies that analyzed microbial dispersal focused on biogeographic distribution patterns or community coalescence based on metacommunity theory at a large scale (Powell *et al.*, 2015; Mansour *et al.*, 2018; Zhang *et al.*, 2018; Brown *et al.*, 2020); even those that included the dispersion of mycorrhizal fungal propagules (Correia *et al.*, 2019; Paz *et al.*, 2021). However, local dispersal limitation has been shown to play a critical role in shaping the distribution and diversity of microorganisms (Telford *et al.*, 2006). Some of these microorganisms, including endospheric fungi, are associated with a host plant with a certain host preference effect (Wagner *et al.*, 2016). The host preference effect results from tight interactions among plants and their associated microbiota leading to the selective recruitment of particular microorganisms. Host plants can thus be considered as the preferential microhabitats of these microorganisms and microbial dispersal limitation may be linked to the distance between host plants (Mony *et al.*, 2020b). In endospheric fungi associated with plant roots, dispersal limitation seems to occur at scales of less than a meter (Mony *et al.*, 2020a), probably because of their low dispersal capacity. Local microbial dispersal among host plants is therefore assumed to be at least partially achieved via the dispersal of spores or hyphal fragments by soil fauna (Lilleskov & Bruns, 2005; Vašutová *et al.*, 2019), by the dispersal of stolons or rhizomes (Vannier *et al.*, 2018), or by inoculation of roots through contact with neighboring plant roots (Mony *et al.*, 2021). The effect of corridors provided by connected host plants may thus promote dispersal fluxes of microorganisms among plants. If this is true, it would result in the same corridor-based predictions for plant symbiotic microorganisms as for macroorganisms.

At the beginning of the development of a plant, colonization of microorganisms is mostly driven by a priority effect (Debray *et al.*, 2022). Fungal species in the immediate vicinity colonize hosts sequentially rather than simultaneously with the first species to arrive, possibly preventing further colonization by latecomers as is the case of arbuscular mycorrhiza (Werner & Kiers, 2015). This early microbial colonization process is quite stochastic because it is strongly determined by the composition of local microbial reservoirs, which can refer to a neutral model assuming homogeneous environments and equivalence of species with respect to their requirements (Bell, 2001). The host plants then progressively regulate the colonization of microorganisms through selective recruitment of particular microorganisms. Such root associated microbial recruitment can be achieved by emitting secondary metabolites such as coumarins (Voges *et al.*, 2019) and volatile compounds (Schulz-Bohm *et al.*, 2018), and by rewarding the most beneficial symbionts in the endosphere (Kiers *et al.*, 2011). All these mechanisms are hypothesized to reduce stochasticity in the microbial assembly and to promote more deterministic microbiota. Therefore, the effect of corridors might be more pronounced over time as root-associated microbial communities become more specific to the host plant species.

In this study, we analyzed the effect of corridors on the dynamics of plant root endospheric mycobiota at metric scale. We used carefully designed mesocosm systems to test two experimental treatments: two isolated patches of *Trifolium repens* L. growing in a matrix of *Brachypodium pinnatum* (L.) P. Beauv., one with a corridor of *T. repens* and one without (Fig.1). After checking that *B. pinnatum* and *T. repens* contained distinct microbiota, we characterized root endospheric mycobiota using high-throughput amplicon sequencing for *T. repens* in connected or isolated patches and analyzed sequence cluster composition, richness and dissimilarity under each treatment. In theory, in a community of sympatric organisms sharing the same habitat, species are neutral if they are irrelevant for their respective successes. In this case, changes in the community composition are expected to be random. In addition to this neutral vision, species can interact with one another leading to determinism in the community assembly process. In natural communities, both neutral and deterministic processes are likely take place and can be partitioned. In the present study, we used neutral community models (NCM) (Sloan *et al.*, 2006) to predict whether the connected patches would facilitate species fluxes between the patches, thereby promoting species coexistence. Based on the predictions of the corridor and the metacommunity theory, we made the three following specific predictions: 1) Connected host plants display higher root endospheric mycobiota species richness than isolated host plants; 2) The root endospheric mycobiota of connected host plants are more similar than that of isolated host plants (i.e. lower beta diversity); 3) Root endospheric mycobiota of connected host plants are less subject to stochasticity than the mycobiota of isolated host plants. These predictions are expected to become more pronounced over time with the development of host plants, because of the reduced importance of the stochastic colonization of roots of prior species, compared with deterministic selection by plants in the assembly of root-associated mycobiota.

**Fig. 1.**
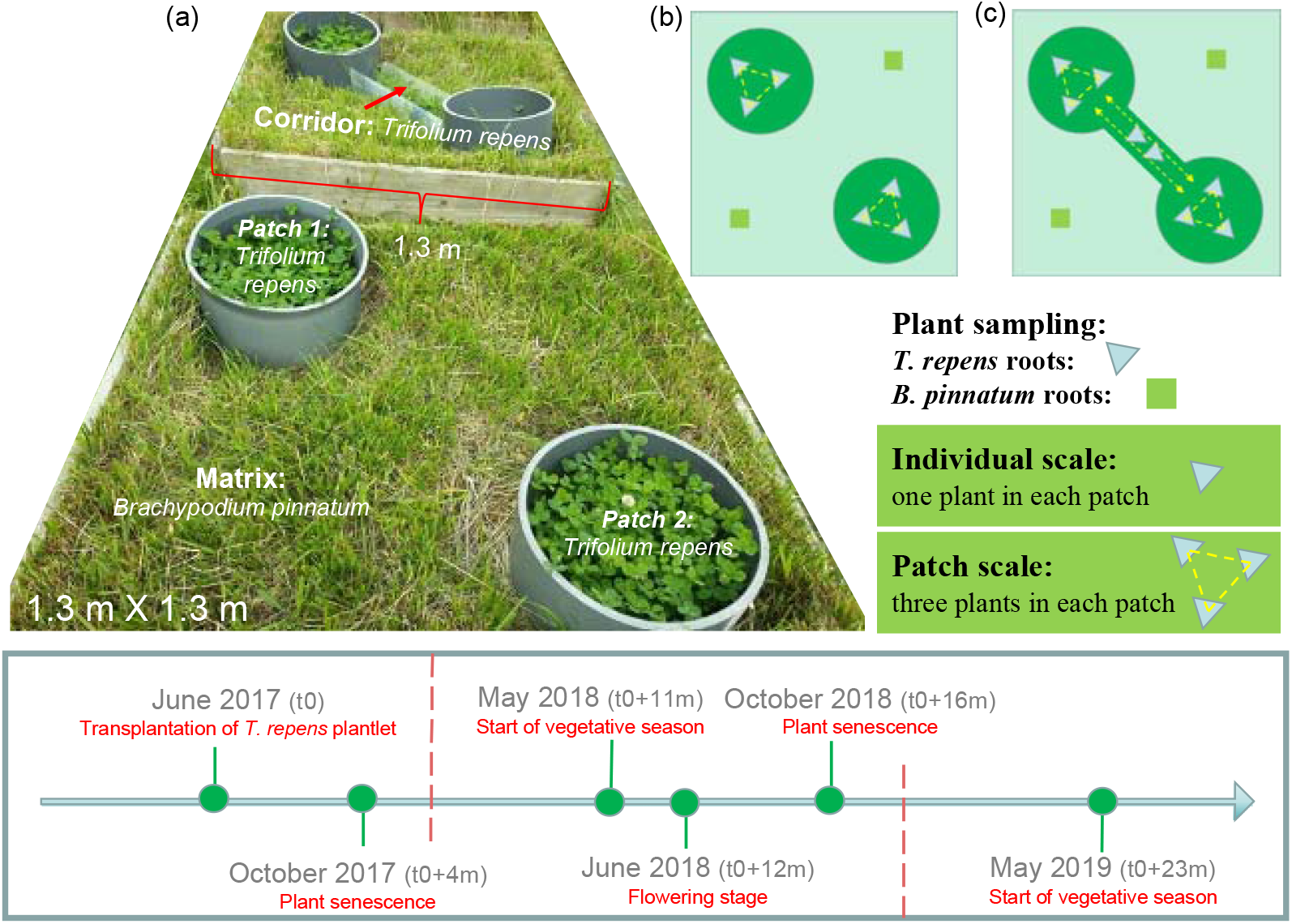
Experimental design of a patch-matrix mesocosm. (a) Photograph of the designed mesocosms used in this study, two patches (D:0.40; H:0.15 m) of *T. repens* were embedded in a matrix (L:1.30 m × W:1.30 m × H:0.25 m) of *B. pinnatum*; in the treatment including a corridor, *T. repens* patches were connected by a narrow corridor (L:0.90 m × W:0.15 m × H:0.15 m) of *T. repens*; (b) plant sampling design for mesocosms with no connecting corridor; (c) plant sampling design for mesocosms with a connecting corridor. Triangles denote *T. repens* roots, rectangles denote *B. pinnatum* roots. Yellow dashed lines indicate potential belowground microbial interactions through plants. Data analyses were conducted at individual and patch scale. At individual scale: the fungal structure of each individual plant was analyzed at each sampling time point. At patch scale: the fungal structure of three *T. repens* plants in the same *T. repens* patch were pooled at each sampling time point (total sequence clusters were pooled with their mean abundances using the R function *aggregate*). In all, 10 mesocosms with a corridor and 9 mesocosms with no corridor were created. The experiment was launched with the transplantation of *T. repens* plantlets in patches with or without a corridor in June 2017 (the 1st year), with five sampling time points: October 2017 (t0+4 months), May 2018 (t0+11 months), June 2018 (t0+12 months), October 2018 (t0+16 months) and May 2019 (t0+23 months), respectively. The corresponding growth stages of *T. repens* are indicated in red below each sampling time point. The red dashed lines denote the end of each year.

## Materials and methods

### Experimental design and plant root sampling

The experimental design consisted of 20 mesocosms located in the common garden of the University of Rennes 1 (France). We tested two treatments, one with and one without a corridor. In both treatments, we planted two patches of *T. repens* (Fabaceae) in a matrix of *B. pinnatum* (Poaceae) (Fig.1A) both being perennial plants. In the corridor treatment, *T. repens* patches were connected by a narrow corridor planted with *T. repens*. *T. repens* and *B. pinnatum* are phylogenetically different and were hence assumed to associate with distinct root microbial assemblages. The experimental treatments thus corresponded to a patch-matrix mesocosm design. We used 10 mesocosms per treatment and these were all set up at the same time (June 9, 2017). In the treatment with no corridor, one mesocosm was lost due to insufficient growth of *T. repens* in the patches. Sampling was performed at five time points (October 2017, May 2018, June 2018, October 2018 and May 2019, Fig.1). At each sampling time point, three ramets of *T. repens* were randomly collected in each of the two patches from all mesocosms, and two ramets were collected in the corridor of mesocosms containing corridors giving a total of 670 samples of *T. repens*. Two ramets of *B. pinnatum* were also collected randomly in the matrix at least 20 cm from the edge of the corridors and at least 10 cm from the edge of the mesocosm giving a total number of 190 samples of *B. pinnatum*. More detailed information on the experimental design, and on cleaning and preparation of the roots is available in the Supporting Materials.

### DNA extraction, amplicon sequencing and bioinformatics

The total DNA of the root associated microbiome was extracted from 100 mg of fresh root powder using a standard protocol with magnetic beads (Sbeadex mini plant kit, LGC Genomics) and an automated protocol (oKtopure robot platform, LGC Genomics) on the Gentyane platform. DNA concentrations were normalized to 12.5 ng/μL after a fluorometric Hoechst DNA assay for subsequent next-generation sequencing. A specific 550 bp of the fungal small subunit (SSU) rRNA gene fragment, including the V4 and V5 regions, was amplified from 50 ng of extracted total DNA of the root-associated microbiome using the primers SSU0817 and NS22B, which conserve 18S rRNA better, thereby limiting overestimation of fungal diversity compared to ITS (e.g. for instance, one given individual Basidiomycota as *Paxillus involutus* or a single spore of a Glomeromycotina, can both display different ITS sequences) but with a disadvantage for the taxonomic resolution. A negative control (i.e. DNA & subsequent PCR products) was also sequenced to detect possible cross-contamination. Both the sequencing library and mass sequencing were performed on the EcogenO platform (https://osur.univ-rennes1.fr/ecogeno). Sequence data were analyzed using the FROGS pipeline (Escudié *et al.*, 2018). Within FROGS, the sequence clustering was performed using SWARM (Mahé *et al.*, 2014) enabling the production of sequence clusters that were not defined by a threshold, and hence close to ASVs or zOTUs, but with the advantage of avoiding artificial overestimation of sequence-based diversity estimates. Sequence clustering was combined with a rigorous chimera removal step (Escudié *et al.*, 2018) followed by the removal of all sequence clusters detected in less than three independent samples and with a threshold of 0.005% of reads. The PhymycoDB database (Mahé *et al.*, 2012) was used for the fungal 18S rRNA gene sequence affiliation. Sequence clusters were filtered using the quality of the affiliations with a threshold of at least 95% BLAST identity and 95% coverage. The number of sequences per sample was normalized to 4 292 sequences by the lowest sequence depth (Fig. S1). This contingency table was used for all subsequent statistical analyses. More detailed information is available in Supporting Materials.

### Statistical analysis

#### Root endospheric mycobiota diversity indices

Root mycobiota diversity was indicated by sequence cluster richness and evenness for the entire community (hereafter termed “global fungi”), and in each phylum group, e.g., Ascomycota, Basidiomycota, Chytridiomycota, Glomeromycota, and Zygomycota, at individual and patch scale (e.g. population scale). At the patch scale, fungal sequence cluster richness and evenness were calculated based on the fungal structure pooled over all *T. repens* samples in the same experimental patch at each sampling time point (the abundance of each sequence cluster was calculated as the mean value of the three ramets sampled in the same patch at that sampling time point). This was calculated using the function *aggregate* in R package stats. Sequence cluster richness was calculated as the total number of sequence clusters present in the whole community of root mycobiota. Evenness was assessed using Pielou’s evenness index based on the original abundance at individual scale and on the mean abundance of sequence clusters at patch scale (Pielou, 1966). Both sequence cluster richness and evenness were calculated using the R package vegan (Oksanen et al., 2022).

To test whether the presence of a corridor, time, and their interactions affected the root mycobiota diversity of global fungi and of each fungal phylum, linear mixed-effects models (LMMs) or generalized linear mixed-effects models (GLMMs) (Bates *et al.*, 2015) for individual and patch scales were used with the presence of a corridor (with versus without), time (5 sampling time points) and their interactions as fixed effects and mesocosm and patch (for individual scale analysis) and mesocosm (for patch scale analysis) as random factors, respectively. We used LMMs with the *lmer* function in the R package lme4 for evenness indices and GLMMs with a negative binomial distribution with the *glmer.nb* function in the R package lme4 for diversity indices, which is appropriate to count data and yielded GLMMs without overdispersion (Zuur *et al.*, 2009). For all models, we tested the model including all the variables. We analyzed the significance of each variable within the model with ANOVA type II sums of squares. We calculated the proportion of variance explained by the fixed effect (marginal R^2^) and by random effects (conditional R^2^) for all analyses (Nakagawa & Schielzeth, 2013). We graphically checked for homoscedasticity, independence and normality of residuals in the tested models. When the models were significant, group comparisons were tested using the *lsmeans* function in the R package lmerTest (Kuznetsova *et al.*, 2017).

#### Root endospheric mycobiota beta-diversity

Fungal beta-diversity was assessed using the dissimilarity index and multivariate analysis. We computed the Bray-Curtis distance matrices using the *vegdist* function in the R package vegan and then visualized them in principal coordinate analysis (PCoA) figures. The effect of the presence of a corridor, time, and their interactions on the structure of root mycobiota was tested with PERMANOVA using the *adonis* function in the R package vegan (Table S1). Because of significant interactive effects of the presence of a corridor and sampling time points, the effect of the presence of a corridor on root endospheric mycobiota composition at individual and patch scales was tested at each sampling time point (Table 1).

**Table 1.** Effect of the presence of a corridor on root endospheric mycobiota structure of *T. repens* at each sampling time point at individual and patch scales. The effects were tested using a PERMANOVA analysis with the *adonis* function in R. Significant results (P < 0.05) are shown in bold.

| Different scales/<br>Sampling time points | Root mycobiota composition<br>at individual scale |  | Root mycobiota<br>composition at patch scale |  |
| --- | --- | --- | --- | --- |
|  | F | P | F | P |
| October 2017 | 1.53 | 0.08 | 1.28 | 0.20 |
| May 2018 | 1.80 | **0.02** | 0.81 | 0.68 |
| June 2018 | 2.21 | **0.01** | 1.23 | 0.24 |
| October 2018 | 2.42 | **0.01** | 2.20 | **0.01** |
| May 2019 | 1.36 | 0.16 | 3.65 | **0.002** |

Bray-Curtis dissimilarity between paired root mycobiota communities of *T. repens* in the same mesocosm at individual scale (all nine possible pairs between three individuals in patch 1 and three individuals in patch 2 of the same mesocosm) and patch scale (only one pair of the same mesocosm) was calculated using the *vegdist* function in the R package vegan. Individual scale corridor effects and time effects on Bray-Curtis distances between root mycobiota of pairwise patches in the same mesocosm were tested using mixed linear models with the mesocosm as a random effect. At patch scale, corridor effects and time effects were tested using t-tests and Tukey honest significant differences respectively.

#### Neutral community model

To detect the assembly process of the root mycobiota in the treatments with and without a corridor, we used a neutral community model proposed by Sloan (Sloan *et al.*, 2006) to predict the relationships between the detection frequency of detection sequence clusters and their relative abundance in a set of local communities (the community in either the treatment with a corridor or without a corridor) across the wider metacommunity (the community of both treatments, i.e. with a corridor and without a corridor). We used the pooled dataset as source data to calculate the null model: there was thus consequently a source data that differed among sampling time points to account for the possible change over time in the pool available for recruitment over time. The sequence clusters from each dataset were subsequently separated into three partitions depending on whether they occurred more frequently than (above the partition), less frequently than (below the partition), or within (neutral partition) the 95% confidence interval of the NCM predictions (Fig. S2). In this model, Nm is an estimate of the dispersal of root mycobiota communities between patches with or without the presence of a corridor. The parameter Nm determines the correlation between the frequency of occurrence and regional relative abundance, where N represents the size of the metacommunity and m the immigration rate. The parameter R^2^ represents the overall fit to the neutral model. Detailed statistics for all the models are provided in Table S2. The 95% confidence intervals around all fitting statistics were calculated by bootstrapping with 1 000 bootstrap replicates.

### Functional prediction of root endospheric mycobiota

To test for the effects of a corridor on fungal groups with particular ecological functions, we combined information from the following databases FUNGuild (Nguyen *et al.*, 2016), FUN^FUN^ version 0.0.3 (Zanne *et al.*, 2020) and FungalTraits version 1.2 (Põlme *et al.*, 2020) to parse fungal sequence clusters with an ecological guild or traits. First, we used FUNGuild.py script in the Python 3 environment to assign the functions of fungi by uploading our own file of taxa to FUNGuild database, then we manually added the complementary information about fungal traits and functions from databases FUN^FUN^ version 0.0.3 and FungalTraits version 1.2. An ecological guild is an index with the potential to indicate the functions of fungal species, but it is important to note that the assignment by the above databases to an ecological guild is currently mostly at the genus taxonomic level. In this study, we focused on three guilds as the main focus in this study: arbuscular mycorrhizae, plant pathogens and saprotrophs. The effect of the presence of a corridor, time, and their interactions on sequence cluster richness and on the abundance (cumulative reads of sequence clusters) of each guild was analyzed using GLMMs with a negative binomial distribution with the *glmer.nb* function in the R package lme4. Group comparisons of FUNGuild sequence cluster richness and abundance at the five sampling time points in treatments with or without a corridor were compared using the *lsmeans* function in the R package lmerTest.

## Results

### Plant species affected the structure of their associated root endospheric mycobiota

The dataset contained a total of 266 fungal sequence clusters dominated by Ascomycota, Basidiomycota, Chytridiomycota, Glomeromycota and Zygomycota (Fig.S3a-b). Even though *B. pinnatum* and *T. repens* shared most of their root endospheric mycobiota (Fig.S4), the abundance of those shared root endospheric mycobiota differed in these two plant species, the root mycobiota community composition of *B. pinnatum* that made up the matrix differed from that of *T. repens* (Fig.2a) across all sampling time points (Fig.S5), demonstrating that these two phylogenetically distant plant species are associated with specific fungal assemblages. The root endospheric mycobiota of *T. repens* was similar in the patches and corridors at each of the five sampling time points (Fig.S6). Despite some overlap, the structure of root-associated mycobiota of *T. repens* differed significantly among the five sampling time points at both individual scale (Fig.2b), and patch scale (Fig.2c). This result indicates that the composition of the mycobiota assemblages changed significantly between the beginning and the end of the 2-year experiment. The composition of the mycobiota depended on the presence of a corridor (only at the individual scale), on the sampling time point and on their interaction both at the individual scale (Table S1 showed the significant effect of the presence of a corridor at all sampling time points except May 2019) and at the patch scale (Table S1 showed significant effect of the presence of a corridor in October 2018 and May 2019.

**Fig. 2.**
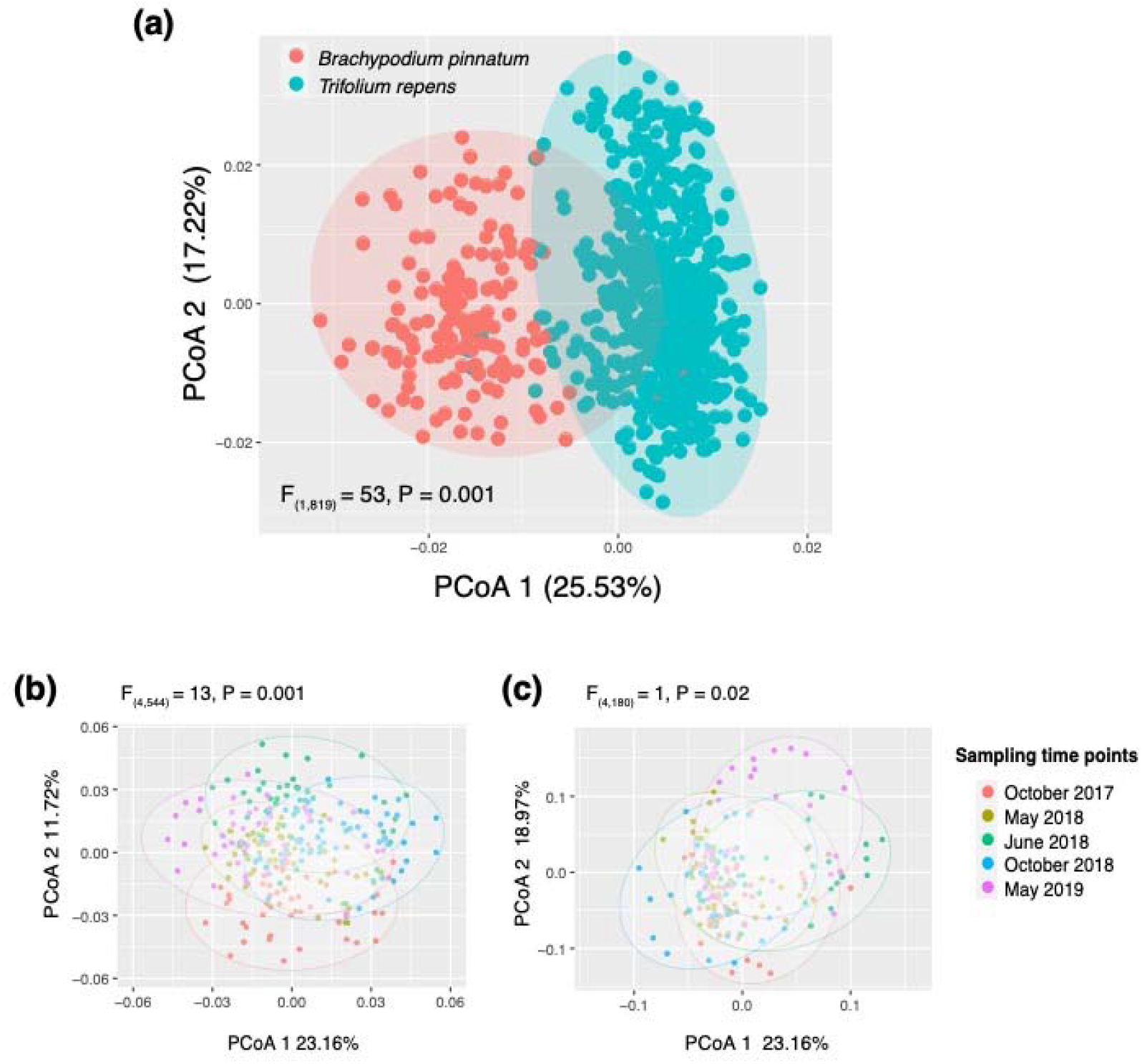
Root endospheric mycobiota structure of *T. repens* and *B. pinnatum* visualized by Principal Coordinate Analysis (PCoA). (a) root mycobiota community composition of both *T. repens* and *Brachypodium pinnatum* at all sampling time points; (b) root mycobiota community composition of only *T. repens* in patches along with five sampling times points at individual scale; (c) root mycobiota community composition of only *T. repens* in patches along with five sampling time points at patch scale. One dot corresponds to the root mycobiota of one *T. repens* sample in panels (a) and (b), one dot corresponds to the root mycobiota of three *T. repens* samples from the same patch in panel (c). Ellipses in each figure represent the 95% confidence interval. Statistics indicated with F and P values obtained from a PERMANOVA analysis with the *adonis* function in R, (a) used the two plant species as factors, (b) and (c) used five sampling time points as factors.

### Ecological corridors promoted plant root endospheric mycobiota diversity

At individual scale, fungal assemblages of connected *T. repens* individuals displayed higher richness than isolated individuals, even though the positive effect of the presence of a corridor on sequence cluster richness was only perceptible in June 2018 and October 2018 (i.e. 12 and 16 months after the start of the experiment) (Fig.3a). At the individual scale, sequence cluster richness depended on the presence of corridors for all phyla at least at one sampling time point (Fig.S7a-e). Higher sequence cluster richness was detected in connected *T. repens* than in isolated *T. repens* in Ascomycota (June 2018, October 2018 and May 2019), Chytridiomycota (October 2018), Zygomycota (June and October 2018). However, and unlike the other phyla, we observed lower sequence cluster richness in Basidiomycota (October 2017), and in Glomeromycota (May 2019) in connected *T. repens* plants than in isolated ones (Fig.S7d-e). Evenness was higher in June 2018 and lower in October 2018 in connected *T. repens* (Fig.3b), with changes only detected in the Chytridiomycota phyla in October 2018 (Fig.S7m). At patch scale, the positive effect of the presence of a corridor on root mycobiota sequence cluster richness was only significant in October 2018 (Fig.3c), while no effect was detected on evenness (Fig.3d). An increase in richness was found in all phyla except Glomeromycota in plants growing in connected patches in October 2018 (Fig.S7f-j).

**Fig. 3.**
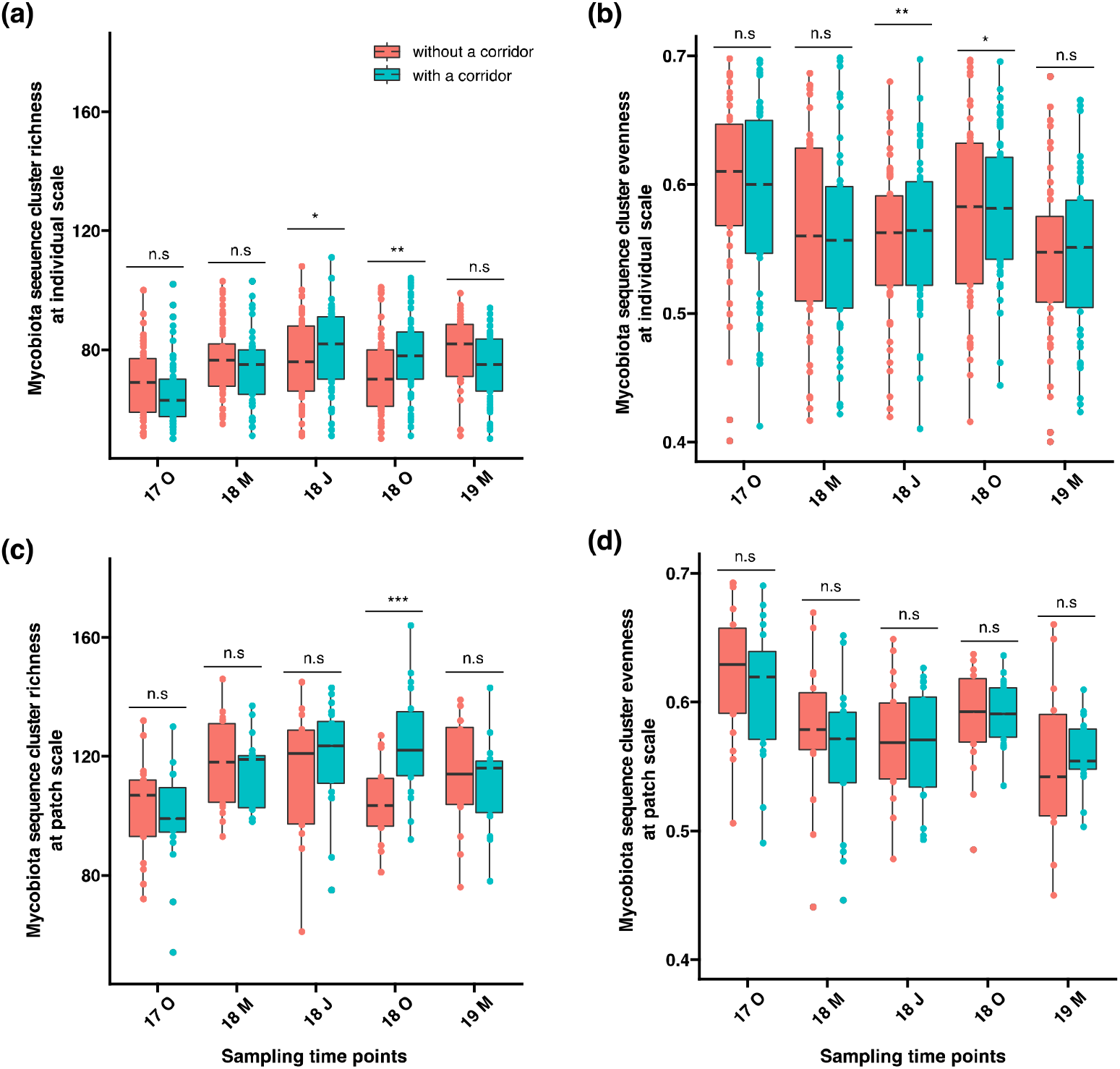
Root endospheric mycobiota diversity for “all fungi” associated with *T. repens* in treatments with and without a corridor over five sampling campaigns at individual and patch scales. Root endospheric mycobiota diversity was calculated as sequence cluster richness and Pielou’s evenness. Indices were calculated and tested at two different biological scales: the individual plant (i.e. one root sample; upper panels a & b) and the patch (i.e. group of three *T. repens* root samples pooled from the same patch; lower panels c & d). (a) and (c) sequence cluster richness of root endospheric mycobiota at individual and patch scales, respectively; (b) and (d) Pielou’s evenness of root endospheric mycobiota at individual and patch scales, respectively. 17 O: October 2017; 18 M: May 2018; 18 J: June 2018; 18 O: October 2018; 19 M: May 2019. Asterisks indicate the significance level of the presence of a corridor on root endospheric mycobiota diversity: * 0.01 < P < 0.05; ** 0.001 < P < 0.01; *** P < 0.001; n.s.: not significant.

### Ecological corridors homogenized plant root endospheric mycobiota composition

The effect of the presence of a corridor on the similarity of *T. repens* root mycobiota composition between patches was assessed using the Bray-Curtis dissimilarity index at the individual and patch scale. While there was no significant effect of the presence of a corridor in October 17 and May 18, the composition of *T. repens* root endospheric mycobiota was more similar in connected individuals and in patches than in isolated individuals and patches at the other sampling time points (June 18, October 18 and May 19) (Fig.4; Fig.S8). In the mesocosms without corridors, the Bray-Curtis dissimilarity index remained stable over time (Fig. 4), whereas it decreased significantly over time in the presence of a corridor (Fig.4). Connected patches showed lower Bray-Curtis dissimilarity in Ascomycota only at individual scale (May 2018, June 2018 and October 2018), in Basidiomycota only at individual scale in June 2018, in Chytridiomycota at both individual (June 2018, October 2018 and May 2019) and patch scales (May 2019) (Fig.S9c and h), in Glomeromycota at both individual (June 2018) and patch scales (June 2018 and October 2018) (Fig.S9d and i), and in Zygomycota only at individual scale in May 2019 (Fig.S9e).

**Fig. 4.**
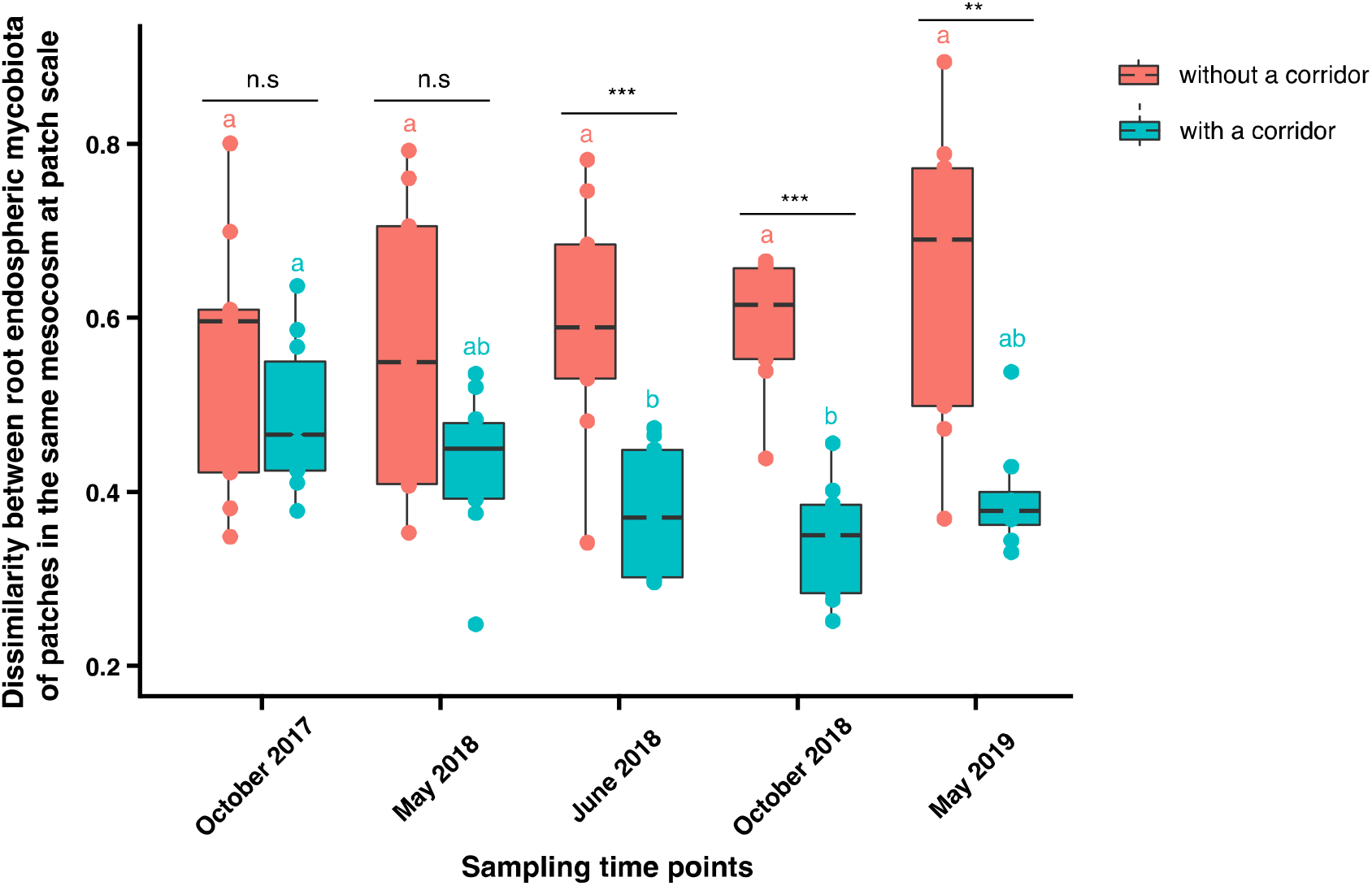
Effect of the presence of a corridor on *T. repens* root endospheric mycobiota dissimilarity at patch scale. *T. repens* root endospheric mycobiota dissimilarity was calculated as Bray-Curtis dissimilarity between the root mycobiota of pairwise *T. repens* patches (i.e. at patch scale) for the same mesocosms with and without a corridor and for the five sampling time points. Pairwise comparisons were conducted on Bray-Curtis dissimilarity between treatments with a corridor and without a corridor using a t-test, and the significance is indicated by asterisks at the top of the bar plots: * 0.01 < P < 0.05; ** 0.01 < P < 0.001; *** P < 0.001; n.s.: not significant. Multiple group comparisons were conducted on Bray-Curtis dissimilarity across sampling campaigns with and without corridors, and the significance is indicated by lowercase letters.

### Ecological corridors boosted the deterministic assembly of plant root endospheric mycobiota

To assess the relative contribution of stochastic and deterministic assembly processes of the root mycobiota, we applied the Sloan neutral model to root endospheric mycobiota sequence data (Sloan *et al.*, 2006) (described in the Method section; Fig.S2). The rationale is to detect more underrepresented and overrepresented sequence clusters than expected in the neutral model and in the calculated neutrality envelope. Both underrepresented and overrepresented sequence clusters, outside the neutrality envelope are assumed to be actively filtered or promoted within the root mycobiota and hence to correspond to deterministic processes of assembly.

Our results may suggest that the stochastic assembly process predominated over the deterministic assembly process, (i.e. measured stochasticity in the richness and abundance of fungal sequence clusters of 45.8%-50.6% and 52.5%-77.6%, respectively) (Fig.5a and Fig.S10a). The estimated stochastic assembly process was higher for endospheric mycobiota of plants growing in patches with no corridor, with higher percentage richness of stochastic sequence clusters recorded at all sampling time points in plants growing in patches with no corridor (50.7%-56.7%) than in patches with a corridor (45.8%-50.6%) (Fig.5a). A higher proportion of sequence clusters (calculated based on their abundance) showing stochastic behavior was detected at two out of the five sampling time points (October 2017 and October 2018) in plants growing in patches with no corridor (Fig.S10a). This observation is likely linked with the higher richness and abundance of neutral sequence clusters detected in patches without a corridor (Fig.5, FigS10 and Fig.S11). Overrepresented sequence clusters dominated in plants growing in patches with a corridor (Fig.5b), potentially due to improved maintenance of biodiversity in patches via dispersal and by limiting the effect of drift. Overrepresented sequence clusters were more frequent in relatively rare species except in October 2017 (Fig.5d, Fig.S10b). Underrepresented sequence clusters were more frequent in plants growing in isolated patches both in richness (Fig.5c, May 2018, June 2018 and May 2019) and in abundance (Fig.S10c, at all sampling time points except October 2017) suggesting greater drift. The presence of a corridor increased ‘Nm’ (Table.S2), where N is the size of the metacommunity and m is the immigration rate, thereby demonstrating the role of the corridor in fungal dispersal. Most neutral sequence clusters in patches both with and without a corridor were assigned to Ascomycota. The highest proportion of underrepresented sequence clusters in plants growing in patches with no corridor was assigned to Basidiomycota at all sampling time points except May 2019 (Fig.S12a, c, e, g and i), while in plants growing in patches with corridors, the highest proportion was assigned to different phyla depending on the sampling time points (Fig.S12b, d, f and h). The highest proportion of overrepresented sequence clusters shifted from Ascomycota to Zygomycota and Basidiomycota depending on the sampling time points for plants growing in isolated patches, and were assigned to Ascomycota and to a lesser extent to Zygomycota (particularly in May and October 2018) in plants growing with in patches with a corridor (Fig.S12b, d, f, h and j). At a higher taxonomic resolution (i.e. species level), *Mycoemilia scoparia* (Zygomycota), *Rhizidium endodporangiatum* (Chytridiomycota), *Rozella allomycis* (Cryptomycota) and *Triparticalcar arcticum* (Chytridiomycota) were the most overrepresented root endospheric sequence clusters with corridor connections detected from the neutral community models (Fig.5d), they were presented as overrepresented sequence clusters in the neutral community models of 5, 4, 3 and 3 sampling time points, respectively. While *Agaricus bisporus* (Basidiomycota), *Anomoporia bombycina* (Basidiomycota), *Geopyxis majalis* (Ascomycota), and *Ambispora fennica* (Glomeromycotina) were the most underrepresented root endospheric sequence clusters without corridor connections (Fig.5d), they were presented as underrepresented sequence clusters in the neutral community models of respectively, 5, 5, 4 and 4 sampling time points.

**Fig. 5.**
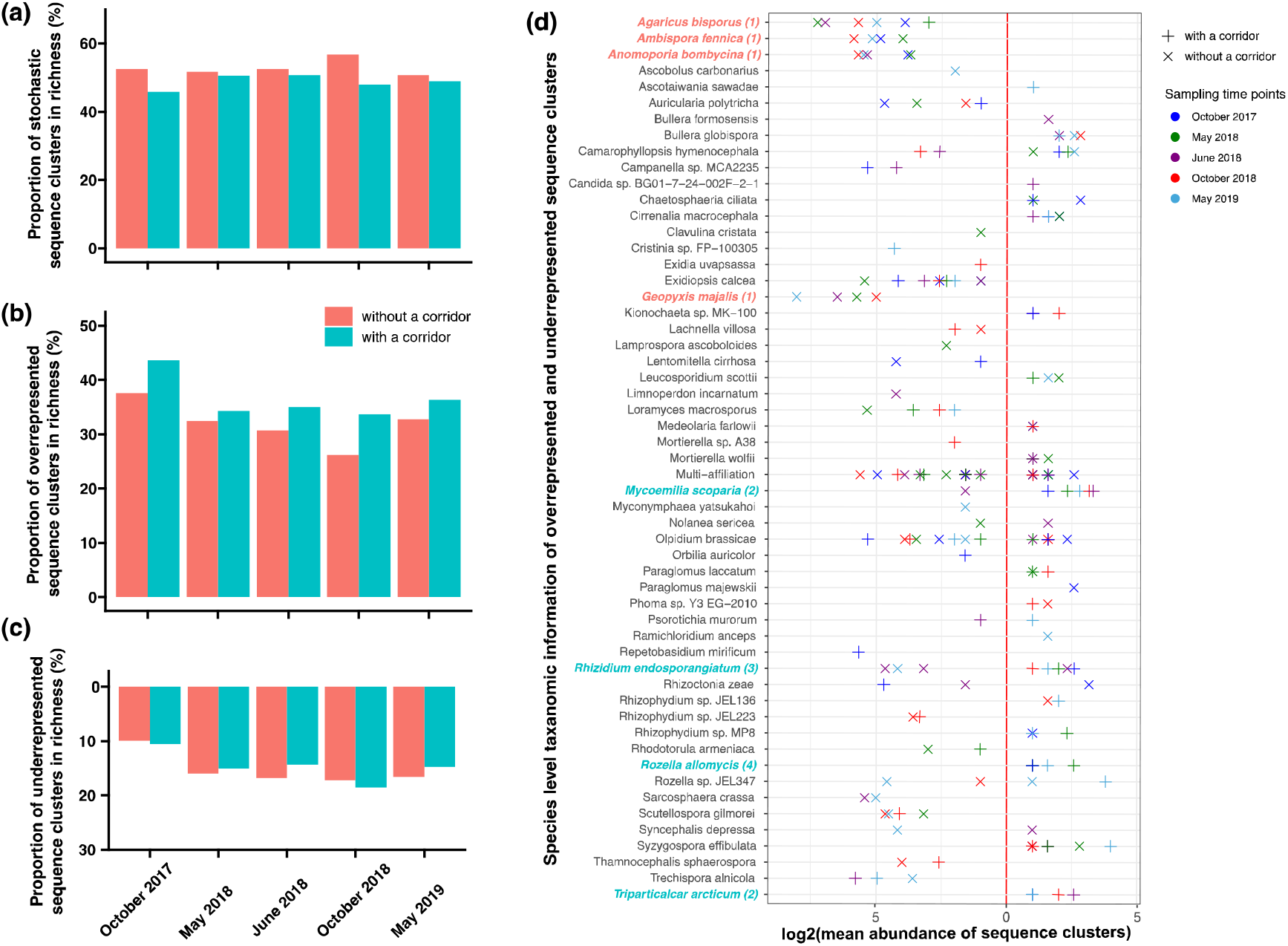
Summary of root endospheric mycobiota assembly patterns with and without corridors at five sampling time points. (a) Proportion of neutral sequence clusters among all the sequence clusters based on their richness in both treatments, i.e. with and without a corridor; (b) proportion of overrepresented sequence clusters among all sequence clusters based on their richness in both treatments, i.e. with and without a corridor; (c) proportion of underrepresented sequence clusters among all the sequence clusters based on their richness in both treatments, i.e. with and without a corridor; (d) species level taxonomic information of overrepresented and underrepresented sequence clusters. In panel (d), underrepresented sequence clusters are shown on the left of the vertical red dashed line. Overrepresented sequence clusters are shown on the right of the vertical red dashed line; the symbol “+” indicates fungal species belonging to treatments with a corridor, the symbol “x” indicates fungal species belonging to treatments without a corridor, symbol colors correspond to the different sampling timepoints indicated in the figure legend on the right. The colors of names of the fungi on the y-axis in panel d correspond to treatments with or without a corridor in panels a-c, the numbers in parentheses after the names of the fungi give the number of sequence clusters belonging to this species.

### Ecological corridors did not promote the spread of plant pathogens

After combining three databases (FUNGuild, FUNFUN and FungalTraits) to predict fungal functions, we were able to assign 138 out of the 266 sequence clusters detected in the *T. repens* root endosphere to ecological guilds, i.e. plant pathogens (16 sequence clusters), arbuscular mycorrhizae (18 sequence clusters) and saprotrophs (77 sequence clusters) (Table S3). Sequence cluster richness and the abundance of plant pathogens did not depend on the presence of a corridor, nor on its interaction with a sampling time point at either the individual or patch scale (Fig.S13a and d, Table 2. Plant pathogen abundance decreased with time both at individual and patch scale (Table 2), but there was no corridor effect on plant pathogen abundance (Fig.S13g and j). Arbuscular mycorrhizal richness decreased with time without a corridor connection at individual scale (Fig.S13b), and arbuscular mycorrhizal richness increased with time with a corridor connection at patch scale (Fig.S13e). Arbuscular mycorrhizal richness was higher with corridor connections in June 2018 at individual scale and in October 2018 at patch scale (Fig.S13b and e). There was no corridor effect on arbuscular mycorrhizal abundance at either individual or patch scale (Fig.S13h and k, Table 2). Saprotroph richness increased with time at both individual and patch scales (Fig.S13c and f, Table 2), while saprotroph abundance decreased with time at both individual and patch scales (Fig.S13i and l, Table 2). Saprotroph richness was higher with corridor connections in June 2018 and October 2018 at individual scale and in October 2018 at patch scale (Fig.S13c and f). There was no corridor effect on saprotroph abundance at either individual or patch scale (Fig.S13i and l, Table 2).

**Table 2.**
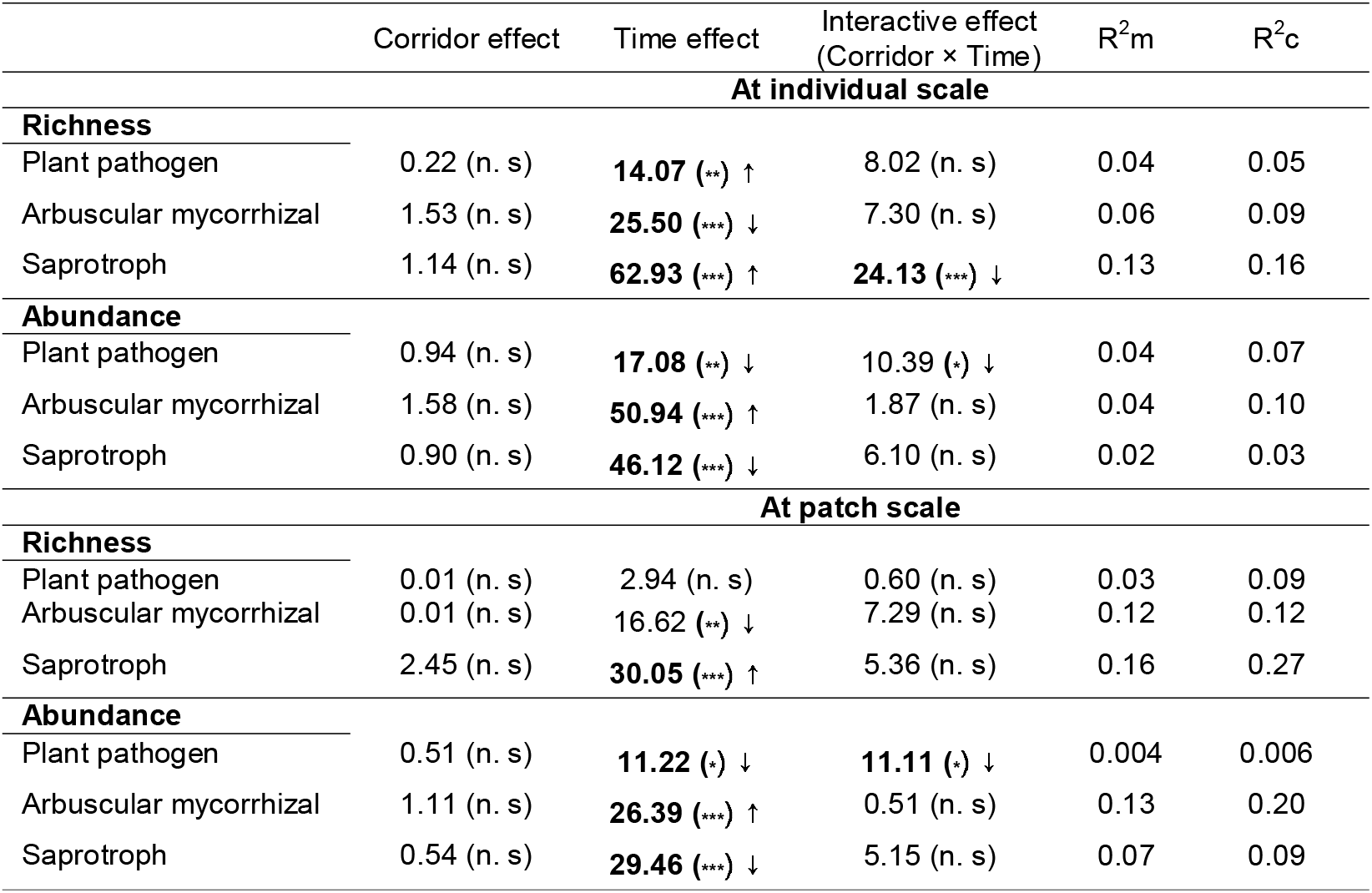
Effects of the presence of a corridor, time, and their interaction on root endospheric mycobiota sequence cluster richness and abundance in each FUNGuild of *T. repens* at individual and patch scales. Random effects of different mesocosms and patches were tested at individual scale, and random effects of different mesocosms were tested at patch scale. R^2^m stands for marginal R squared, and denotes total variance explained by regression, coefficient of determination in the model with only fixed effects; R^2^c stands for conditional R squared, and denotes total variance explained by regression-coefficient of determination in the full random model. Significant results are shown with asterisks in parentheses after the Chisq value: * 0.01 < P < 0.05; ** 0.001 < P < 0.01; *** P < 0.001. n.s.: not significant. Upward and downward arrows denote positive or negative effects of explanatory variables in all the models, respectively. Significant results (P < 0.05) are shown in bold.

## Discussion

### Plant root endospheric mycobiota was shaped by the presence of corridors

While the structure of sequence clusters of *T. repens* roots was marked by strong time dynamics, we demonstrate that plant mycobiota was shaped by connectivity between patches of individual plants. Depending on the sampling time points, the presence of a corridor influenced root endospheric fungal structure, richness and/or evenness, and fungal diversity at all scales from the individual plant, to plant patches and between patches. Our experimental design was based on the assumption that two phylogenetically distant plants such as *B. pinnatum* and *T. repens* were associated with different root fungal communities, and could thus be used to simulate the isolation or connectivity of fungal assemblages associated with *T. repens* individuals. This assumption was confirmed throughout the experiment with contrasted fungal assemblages in the two plant species, but similar assemblages between *T. repens* mycobiota originating from patches or corridors, suggesting a strong host preference effect. Edge effects (Tscharntke *et al.*, 2012) were limited with little species spillover from the *B. pinnatum* matrix to *T. repens* patches despite the likely existence of contacts between the roots of the two plant species. Such a spillover edge effect was also absent in corridors, despite their longer edges and linear forms that result in a higher ratio between the length of edges to the amount of habitat that would render them more prone to the influence of *B. pinnatum* growing outside the corridors. The rooting system of the host plants may provide available niches for fungal species with a certain host preference effect and correspond to particular habitats to colonize, or through which to disperse. Dispersal may involve not only a given organism or species, but also small or large parts of communities leading to community coalescence (Rillig *et al.*, 2015), i.e. the mixing of different communities, from which homogenization of species composition and community structure would be expected. The expected homogenization influence of corridors was not detected at the very beginning of the experiment but was detected 10 to 12 months later. This time lag response may be explained by the succession process involved in the assembly of fungi associated with plants throughout the course of their development from germination to the mature stage. Indeed, newly developing roots are initially randomly colonized by the microbial species present in the soil where the roots are developing (Dini-Andreote & Raaijmakers, 2018; Hu *et al.*, 2020). This colonization can take place through fungal propagules, hyphal fragments or spores that are present locally in the soil (Smith & Read, 2008). Then during the course of their development, individual plants progressively select among the fungi colonizing the endosphere those that are the most favorable for their own development through active recruitment or filtration processes, such as the carbon rewarding process (Kiers *et al.*, 2011). As we found in the present study, plant filtering modifies species composition over time, with a decrease in fungal species evenness in fungal assemblages associated with individual plants. Our results suggest that in addition to this plant filtering process, the spatial configuration of the plant influences the process of fungal succession. The influence of the host plant configuration on root associated mycobiota has already been demonstrated in *B. pinnatum* plant species (Mony *et al.*, 2020b), especially due to the functional connectivity it provided among plants (Mony *et al.*, 2020a). The present study clearly demonstrates that this connectivity effect really does exist. To validate the assumptions about the dispersal and colonization process, it would be interesting to undertake complementary experiments with tagged fungi that could be surveyed over time in the presence and absence of corridors. This could be done using a range of fungi selected for their contrasted colonization abilities in order to obtain more detailed information to explain the community-level dispersal we addressed here.

### Connected host plants displayed higher root endospheric mycobiota species richness and more similar species structure than isolated host plants

Confirming our first prediction, we demonstrated that the presence of a corridor increased the endospheric fungal sequence cluster richness associated with *T. repens* roots, and that this effect was more pronounced with plant aging and in later phenological stages, notably after the plants have flowered. This effect on fungal species richness was accompanied by a change in species composition that was already detected at the first sampling time point and remained significant right up to the end of the experiment. Changes were observed earlier at the individual scale (in composition in October 2017, in richness in June 2018) than at the patch scale (in composition and richness in October 2018), reflecting a progressive process that reached and characterized the entire population of host plants. In addition, we confirmed our second prediction of a more similar endospheric mycobiota among *T. repens* individuals in connected patches than in isolated ones. This was evidenced by a decrease in root endospheric mycobiota dissimilarity associated with connected *T. repens* individuals compared to isolated ones (detected at both the individual and patch scales). This effect on beta-diversity was detected after June 2018, i.e. one year after the start of the experiment, and until the end of the experiment. This result confirms the fact that, despite starting from a different and stochastic root endospheric mycobiota, corridors tend to homogenize root-associated fungal composition among connected plants, most likely through a long-term effect. Such time-lag effects in response to landscape structure are referred to as a colonization credit and have been repeatedly demonstrated in the case of macroorganisms (Kuussaari *et al.*, 2009). There have only been a few demonstrations of such processes in microorganisms especially in response to habitat connectivity (Mony *et al.*, 2022), probably because of the limited number of studies based on time-series analysis of microbial composition. The present work demonstrates the importance of analyzing temporal dynamics in microbial response to host landscapes.

In landscape ecology, the positive effect of the presence of corridors on diversity in the landscape is often explained by two main processes: facilitated dispersal among habitat patches (Rosenberg *et al.*, 1997; Beier & Noss, 1998), and/or an increase in habitat availability, which refers to the habitat amount hypothesis (Fahrig, 2013). Our results suggest that these processes may also apply to fungi, at least at the metric scale considered in this study. Dispersal of mycorrhizal fungi among patches via the experimented corridor can be achieved by root contacts (Smith & Read, 2008), but fungi also disperse through the clonal development of stolons (Vannier *et al.*, 2018, 2019). Additional dispersal processes may involve litter decomposition and a legacy effect on the soil reservoir after the plant leaf senescence stage that influenced the microbial composition of the sampling campaign in October, which was clearly shown to have an obvious seasonal effect. Indeed, litter composition has a strong impact on soil fungal composition (Habtewold *et al.*, 2020). In addition to improved species coexistence, corridors led to species homogenization among patches by promoting preferential dispersal of particular fungi along the corridor, leading to more deterministic assemblages. We confirmed our third prediction that root endospheric mycobiota of *T. repens* in connected patches were less subjected to stochasticity than in isolated *T. repens* patches, suggesting a buffering effect of drift through corridors. Despite an existing fungal host plant preference confirmed herein, the drift effect in isolated patches could result from the priority effect, i.e., the first fungal colonizers and plant filtration processes are expected to lead to divergent root endospheric mycobiota composition, with the strength of the drift effect expected to be positively correlated with the heterogeneity of available fungal propagules among disconnected patches. This is in line with the observation that among the most underrepresented fungi in patches with no corridor, we found an arbuscular mycorrhiza (*Ambispora fennica*, Figure 5d) for which an existing priority effect has already been demonstrated (Werner & Kiers, 2015). The other underrepresented-fungi in the unconnected patches are often known as saprotrophs (Table S3). Similarly, the majority of the overrepresented fungi in the presence of a corridor are also known as saprotrophs. In both cases these fungi colonize the endosphere of healthy *T. repens*, i.e. a similar habitat. This can be interpreted as a consequence of competitive exclusion and competitive behavior for root colonization. More specifically, a competition-colonization trade-off, i.e. the idea that if bad competitors can escape from their competitive superiors through better dispersal and can persist by finding new habitat, this could explain the coexistence of functionally redundant fungi (Smith *et al.*, 2018). Corridors between patches facilitating the dispersal can explain the overrepresented root-colonizing fungal saprotrophs. Fungal competition-colonization trade-offs could be seen as an important hypothesis to be further explored to link ecological functions and traits to community assembly and dynamics, possibly through experiments using fungal synthetic communities to colonize plants.

The importance of habitat availability for microscopic fungal species is questionable, as many processes that define the carrying capacity of the roots of individual plants for fungi are remain unknown. Interactions among microbial components (e.g. fungus-fungus; fungus-bacteria and others) that constitute the plant microbiota might indeed impact the carrying capacity of plants for microorganisms. Interactions include competition for habitat, available nutrients, exclusion through antimicrobial compounds and reciprocally, via facilitation processes such as cross feeding, co-metabolism, evolution of dependencies (Mataigne *et al.*, 2021), but these ideas are still mostly theoretical and experimental/empirical supports are needed. Beyond these microbial interactions that putatively at least partially explain the plant microbiota, we cannot exclude possible indirect effects of *T. repens* on soil characteristics that represent an indirect effect on habitat conditions for fungi. It has long been known that, like other Fabaceae species associated with a Rhizobium, *T. repens* could lead to enriched bioavailable soil nitrogen (Ruschel *et al.*, 1979), leading to changes in habitat characteristics (including abiotic conditions). Changes in soil characteristics, and especially in nitrogen, are known to affect both fungal and bacterial activity in soil (Demoling *et al.*, 2008; Wang *et al.*, 2021), and which also possibly result in a change in root endospheric microbiota (Mareque *et al.*, 2018). Further work is necessary to disentangle the direct effect of *T. repens* corridors on microbial dispersal from changes in soil abiotic conditions resulting in changes in the soil microorganism reservoir.

### The presence of a corridor affected the fungal species that form the plant root endospheric mycobiota differently

Corridors increased fungal richness in all phyla except Glomeromycota, both at individual (except Basidiomycota) and patch scale; and despite a wide range of dispersal strategies among phyla. The absence of an effect on Glomeromycota may result from a less pronounced host preference effect on this phylum, thereby increasing the influence of plants developing in the vicinity (Hausmann & Hawkes, 2009; Mony *et al.*, 2020a). Because the roots of focal and neighboring plants may intermingle, exchange of the entire set of the fungal assemblages might occur between the two habitats illustrating a potential coalescence process among host plants (Rillig *et al.*, 2015). At the last sampling point, the reduced richness in Glomeromycota in patches with a corridor may be related to a strong effect of competitive interactions among fungi belonging to Glomeromycota fungi within roots (Maherali & Klironomos, 2007) driving species assembly. In contrast, arbuscular mycorrhizal richness was increased over time in the presence of a corridor, probably because the disappearance of some Glomeromycota species released ecological niches for the other arbuscular mycorrhizal species. A better understanding of how corridors affect certain components of the plant mycobiota at short spatial scales could be important, considering the wide range of functions fulfilled by fungi for plants (Vandenkoornhuyse *et al.*, 2015). The results we obtained based on functional guilds revealed no significant effect of connectivity on plant pathogens, and only a weak effect on arbuscular mycorrhizae and saprotrophs, however, these results should be interpreted with caution due to the lack of available information about almost half of sequence clusters in our dataset.

### Corridors as a key to understanding plant associated mycobiota assembly: toward a concept of biotic corridors

Through this work, which was based on a simple but robust experimental design with two patches of plants isolated or connected by a linear corridor made of the same plants, we demonstrated that plants also provide biotic corridors for fungi, with effects on the composition, alpha and beta diversity of root associated fungal communities. Our results followed the same predictions as those concerning corridors for macroorganisms (Pardini *et al.*, 2005). An original aspect of this study is that we detected such corridor effects at the community level despite the wide range of modes of dispersal. Community-wide assessment of corridor effects is quite recent in landscape ecology for macroorganisms (Uroy *et al.*, 2019), while it is one of the very first pieces of evidence for microorganisms (see Mony et al., 2022 for a review on habitat corridor studies for microbes). Because such effects are demonstrated at the community level, these results also demonstrate that most fungi are limited to short spatial scales for their dispersal. In addition, ecological corridors are often defined by a habitat (Beier & Noss, 1998; Gilbert-Norton *et al.*, 2010; Fletcher *et al.*, 2016). Here, we demonstrated that the corridors provided by host plants may provide suitable habitats for certain fungi to promote their exchanges between two connected patches, and mitigate the negative influence of isolation on the structure of endospheric mycobiota. The corridors could comprise biotic corridors that differ from the classical corridor concept because of the tight interactions of a host and its microbiota that override the effect of the characteristics of the physical environment (e.g. here habitat is better defined by the host plants than by the abiotic conditions). Such a biotic corridor concept could widely apply to microbes due to the large number of host-associated microbes. Whether the corridor affects dispersal independently of a potential indirect effect of *T. repens* on soil characteristics or not remains an open question. Plants are known to be associated with preferential fungal assemblages, here we demonstrated that, for fungi, a host preference effect can also influence processes at the larger spatial scale of the micro-landscape, thereby promoting connectivity of fungi from one patch of host plants to another. Such an effect could explain the marked spatial heterogeneity of fungi recorded even at small spatial scales (Bahram *et al.*, 2015). This paper illustrates a new concept of biotic corridors for microorganisms that offers opportunities to advance our theoretical understanding of fungal assembly, and for the manipulation of *in situ* mycobiota structure and its application in agriculture through plant configuration.

## Acknowledgements

This work was supported by regional aid for H2020-MSCA-IF projects 2019 Seal of Excellence - CORRiBIOM to J.H., C.M. and P.V.. We thank Remi Bodiguel and Thierry Fontaine from the ECOLEX platform for designing the mesocosms. We thank George A. Kowalchuk from Utrecht University and Ecobio members from University of Rennes 1 for helpful comments during departmental seminars.

## Author contributions

C.M. and P.V. conceived the study and the methodology. C.M., P.V., F.K. collected the data. F.K. and R.C.-V. performed the sequence analyses. J.H. performed statistical analysis of the data and wrote the manuscript with the help of C.M. and P.V. All the authors contributed critically to interpreting the results and to the drafts and gave their final approval for publication.

## Data availability

Sequencing reads from root endospheric mycobiota have been deposited in the Sequence Read Archive (SRA: https://submit.ncbi.nlm.nih.gov) and are accessible under accession number PRJNA793385. Data that are relevant to this manuscript and the scripts used in the computational analyses are available at https://github.com/HuJamie/Microbial.Corridor. All other study data are included in the article and/or in supporting information.

## SUPPORTING INFORMATION

– Supporting Materials

– Supporting references

– Figs. S1 to S13

– Tables S1 to S3

## References

Arévalo JR, Otto R, Escudero C, Fernández-Lugo S, Arteaga M, Delgado JD, Fernández-Palacios JM. 2010. Do anthropogenic corridors homogenize plant communities at a local scale? A case studied in Tenerife (Canary Islands). Plant Ecology 209: 23–35.

Baas Becking, L. G. M. 1934. Geobiologie of Inleiding Tot de Milieukunde. The Hague: W. P. Van Stockum and Zoon.

Bahram M, Peay KG, Tedersoo L. 2015. Local‐scale biogeography and spatiotemporal variability in communities of mycorrhizal fungi. New Phytologist 205: 1454–1463.

Bates D, Mächler M, Bolker B, Walker S. 2015. Fitting Linear Mixed‐Effects Models Using lme4. Journal of Statistical Software 67.

Beier P, Noss RF. 1998. Do Habitat Corridors Provide Connectivity? Conservation Biology 12: 1241–1252.

Bell G. 2001. Neutral Macroecology. Science 293: 2413–2418.

Brown JH, Kodric-Brown A. 1977. Turnover Rates in Insular Biogeography: Effect of Immigration on Extinction. Ecology 58: 445–449.

Brown JJ, Mihaljevic JR, Des Marteaux L, Hrček J. 2020. Metacommunity theory for transmission of heritable symbionts within insect communities. Ecology and Evolution 10: 1703–1721.

Chagnon P, Bradley RL, Klironomos JN. 2020. Mycorrhizal network assembly in a community context: The presence of neighbours matters (P Mariotte, Ed.). Journal of Ecology 108: 366–377.

Correia M, Heleno R, Silva LP da, Costa JM, Rodríguez-Echeverría S. 2019. First evidence for the joint dispersal of mycorrhizal fungi and plant diaspores by birds. New Phytologist 222: 1054–1060.

Damschen EI. 2006. Corridors Increase Plant Species Richness at Large Scales. Science 313: 1284–1286.

Debray R, Herbert RA, Jaffe AL, Crits-Christoph A, Power ME, Koskella B. 2022. Priority effects in microbiome assembly. Nature Reviews Microbiology 20: 109–121.

Demoling F, Ola Nilsson L, Bååth E. 2008. Bacterial and fungal response to nitrogen fertilization in three coniferous forest soils. Soil Biology and Biochemistry 40: 370–379.

Diamond JM. 1976. Island Biogeography and Conservation: Strategy and Limitations. Science 193: 1027–1029.

Dini-Andreote F, Raaijmakers JM. 2018. Embracing Community Ecology in Plant Microbiome Research. Trends in Plant Science 23: 467–469.

Escudié F, Auer L, Bernard M, Mariadassou M, Cauquil L, Vidal K, Maman S, Hernandez-Raquet G, Combes S, Pascal G. 2018. FROGS: Find, Rapidly, OTUs with Galaxy Solution (B Berger, Ed.). Bioinformatics 34: 1287–1294.

Fahrig L. 2013. Rethinking patch size and isolation effects: the habitat amount hypothesis (K Triantis, Ed.). Journal of Biogeography 40: 1649–1663.

Fletcher RJ, Burrell NS, Reichert BE, Vasudev D, Austin JD. 2016. Divergent Perspectives on Landscape Connectivity Reveal Consistent Effects from Genes to Communities. Current Landscape Ecology Reports 1: 67–79.

Gilbert-Norton L, Wilson R, Stevens JR, Beard KH. 2010. A Meta‐Analytic Review of Corridor Effectiveness: Corridor Meta‐Analysis. Conservation Biology 24: 660–668.

Groppe K, Steinger T, Schmid B, Baur B, Boller T. 2001. Effects of habitat fragmentation on choke disease (Epichloë bromicola) in the grass Bromus erectus: fragmentation and choke disease. Journal of Ecology 89: 247–255.

Habtewold JZ, Helgason BL, Yanni SF, Janzen HH, Ellert BH, Gregorich EG. 2020. Litter composition has stronger influence on the structure of soil fungal than bacterial communities. European Journal of Soil Biology 98: 103190.

Haddad NM. 2015. Corridors for people, corridors for nature. Science 350: 1166–1167.

Haddad NM, Brudvig LA, Clobert J, Davies KF, Gonzalez A, Holt RD, Lovejoy TE, Sexton JO, Austin MP, Collins CD, et al. 2015. Habitat fragmentation and its lasting impact on Earth’s ecosystems. Science Advances 1: e1500052.

Haddad NM, Brudvig LA, Damschen EI, Evans DM, Johnson BL, Levey DJ, Orrock JL, Resasco J, Sullivan LL, Tewksbury JJ, et al. 2014. Potential Negative Ecological Effects of Corridors: Negative Effects of Corridors. Conservation Biology 28: 1178–1187.

Hanson CA, Fuhrman JA, Horner-Devine MC, Martiny JBH. 2012. Beyond biogeographic patterns: processes shaping the microbial landscape. Nature Reviews Microbiology 10: 497–506.

Hausmann NT, Hawkes CV. 2009. Plant neighborhood control of arbuscular mycorrhizal community composition. New Phytologist 183: 1188–1200.

Hilty JA, Lidicker WZ, Merenlender AM. 2006. Corridor ecology: the science and practice of linking landscapes for biodiversity conservation. Washington, DC: Island Press.

Hu J, Wei Z, Kowalchuk GA, Xu Y, Shen Q, Jousset A. 2020. Rhizosphere microbiome functional diversity and pathogen invasion resistance build up during plant development. Environmental Microbiology: 1462‐2920.15097.

Jamoneau A, Chabrerie O, Closset-Kopp D, Decocq G. 2012. Fragmentation alters beta‐diversity patterns of habitat specialists within forest metacommunities. Ecography 35: 124–133.

Kiers ET, Duhamel M, Beesetty Y, Mensah JA, Franken O, Verbruggen E, Fellbaum CR, Kowalchuk GA, Hart MM, Bago A, et al. 2011. Reciprocal Rewards Stabilize Cooperation in the Mycorrhizal Symbiosis. Science 333: 880–882.

Kuussaari M, Bommarco R, Heikkinen RK, Helm A, Krauss J, Lindborg R, Öckinger E, Pärtel M, Pino J, Rodà F, et al. 2009. Extinction debt: a challenge for biodiversity conservation. Trends in Ecology & Evolution 24: 564–571.

Kuznetsova A, Brockhoff PB, Christensen RHB. 2017. lmerTest Package: Tests in Linear Mixed Effects Models. Journal of Statistical Software 82.

Langenheder S, Lindström ES. 2019. Factors influencing aquatic and terrestrial bacterial community assembly. Environmental Microbiology Reports 11: 306–315.

Leibold MA, Holyoak M, Mouquet N, Amarasekare P, Chase JM, Hoopes MF, Holt RD, Shurin JB, Law R, Tilman D, et al. 2004. The metacommunity concept: a framework for multi‐scale community ecology: The metacommunity concept. Ecology Letters 7: 601–613.

Lilleskov EA, Bruns TD. 2005. Spore dispersal of a resupinate ectomycorrhizal fungus, Tomentella sublilacina, via soil food webs. Mycologia 97: 762–769.

Mahé S, Duhamel M, Le Calvez T, Guillot L, Sarbu L, Bretaudeau A, Collin O, Dufresne A, Kiers ET, Vandenkoornhuyse P. 2012. PHYMYCO‐DB: A Curated Database for Analyses of Fungal Diversity and Evolution (D Steinke, Ed.). PLoS ONE 7: e43117.

Mahé F, Rognes T, Quince C, de Vargas C, Dunthorn M. 2014. Swarm: robust and fast clustering method for amplicon‐based studies. PeerJ: e593.

Maherali H, Klironomos JN. 2007. Influence of Phylogeny on Fungal Community Assembly and Ecosystem Functioning. Science 316: 1746–1748.

Mansour I, Heppell CM, Ryo M, Rillig MC. 2018. Application of the microbial community coalescence concept to riverine networks: Riverine microbial community coalescence. Biological Reviews 93: 1832–1845.

Mareque C, da Silva TF, Vollú RE, Beracochea M, Seldin L, Battistoni F. 2018. The Endophytic Bacterial Microbiota Associated with Sweet Sorghum (Sorghum bicolor) Is Modulated by the Application of Chemical N Fertilizer to the Field. International Journal of Genomics 2018: 1–10.

Martiny JBH, Bohannan BJM, Brown JH, Colwell RK, Fuhrman JA, Green JL, Horner-Devine MC, Kane M, Krumins JA, Kuske CR, et al. 2006. Microbial biogeography: putting microorganisms on the map. Nature Reviews Microbiology 4: 102–112.

Mataigne V, Vannier N, Vandenkoornhuyse P, Hacquard S. 2021. Microbial Systems Ecology to Understand Cross‐Feeding in Microbiomes. Frontiers in Microbiology 12: 780469.

Miller ET, Svanbäck R, Bohannan BJM. 2018. Microbiomes as Metacommunities: Understanding Host‐ Associated Microbes through Metacommunity Ecology. Trends in Ecology & Evolution 33: 926–935.

Mony C, Brunellière P, Vannier N, Bittebiere A-K, Vandenkoornhuyse P. 2020a. Effect of floristic composition and configuration on plant root mycobiota: a landscape transposition at a small scale. New Phytologist 225: 1777–1787.

Mony C, Gaudu V, Ricono C, Jambon O, Vandenkoornhuyse P. 2021. Plant neighbors shape fungal assemblages associated with plant roots: a new understanding of niche‐partitioning in plant communities. Functional Ecology: 1365‐2435.13804.

Mony C, Uroy L, Khalfallah F, Haddad N, Vandenkoornhuyse P. 2022. Landscape connectivity for the invisibles. Ecography 2022.

Mony C, Vannier N, Brunellière P, Biget M, Coudouel S, Vandenkoornhuyse P. 2020b. The influence of host‐plant connectivity on fungal assemblages in the root microbiota of Brachypodium pinnatum. Ecology 101.

Mouquet N, Loreau M. 2003. Community Patterns in Source‐Sink Metacommunities. The American Naturalist 162: 544–557.

Nakagawa S, Schielzeth H. 2013. A general and simple method for obtaining R from generalized linear mixed‐effects models (RB O’Hara, Ed.). Methods in Ecology and Evolution 4: 133–142.

Nemergut DR, Schmidt SK, Fukami T, O’Neill SP, Bilinski TM, Stanish LF, Knelman JE, Darcy JL, Lynch RC, Wickey P, et al. 2013. Patterns and Processes of Microbial Community Assembly. Microbiology and Molecular Biology Reviews 77: 342–356.

Nguyen NH, Song Z, Bates ST, Branco S, Tedersoo L, Menke J, Schilling JS, Kennedy PG. 2016. FUNGuild: An open annotation tool for parsing fungal community datasets by ecological guild. Fungal Ecology 20: 241–248.

Oksanen et al. 2022. Vegan: Community Ecology Package. R package Version 2.6‐2.

Pardini R, de Souza SM, Braga-Neto R, Metzger JP. 2005. The role of forest structure, fragment size and corridors in maintaining small mammal abundance and diversity in an Atlantic forest landscape. Biological Conservation 124: 253–266.

Paz C, Öpik M, Bulascoschi L, Bueno CG, Galetti M. 2021. Dispersal of Arbuscular Mycorrhizal Fungi: Evidence and Insights for Ecological Studies. Microbial Ecology 81: 283–292.

Pielou EC. 1966. The measurement of diversity in different types of biological collections. Journal of Theoretical Biology 13: 131–144.

Põlme S, Abarenkov K, Henrik Nilsson R, Lindahl BD, Clemmensen KE, Kauserud H, Nguyen N, Kjøller R, Bates ST, Baldrian P, et al. 2020. FungalTraits: a user‐friendly traits database of fungi and fungus‐like stramenopiles. Fungal Diversity 105: 1–16.

Powell JR, Karunaratne S, Campbell CD, Yao H, Robinson L, Singh BK. 2015. Deterministic processes vary during community assembly for ecologically dissimilar taxa. Nature Communications 6: 8444.

Rillig MC, Antonovics J, Caruso T, Lehmann A, Powell JR, Veresoglou SD, Verbruggen E. 2015. Interchange of entire communities: microbial community coalescence. Trends in Ecology & Evolution 30: 470–476.

Rosenberg DK, Noon BR, Meslow EC. 1997. Biological Corridors: Form, Function, and Efficacy. BioScience 47: 677–687.

Ruschel AP, Salati, E., Vose, P. B. 1979. Nitrogen Enrichment of soil and Plant by Rhizobium Phaseoli-Phaseolus Vulgaris Symbiosis. 51(3):425–429.

Schulz-Bohm K, Gerards S, Hundscheid M, Melenhorst J, de Boer W, Garbeva P. 2018. Calling from distance: attraction of soil bacteria by plant root volatiles. The ISME Journal 12: 1252–1262.

Sloan WT, Lunn M, Woodcock S, Head IM, Nee S, Curtis TP. 2006. Quantifying the roles of immigration and chance in shaping prokaryote community structure. Environmental Microbiology 8: 732–740.

Smith SE, Read DJ. 2008. Mycorrhizal symbiosis. Amsterdam Boston: Academic Press.

Smith GR, Steidinger BS, Bruns TD, Peay KG. 2018. Competition–colonization tradeoffs structure fungal diversity. The ISME Journal 12: 1758–1767.

Telford RJ, Vandvik V, Birks HJB. 2006. Dispersal Limitations Matter for Microbial Morphospecies. Science 312: 1015–1015.

Tewksbury JJ, Levey DJ, Haddad NM, Sargent S, Orrock JL, Weldon A, Danielson BJ, Brinkerhoff J, Damschen EI, Townsend P. 2002. Corridors affect plants, animals, and their interactions in fragmented landscapes. Proceedings of the National Academy of Sciences 99: 12923–12926.

Tscharntke T, Tylianakis JM, Rand TA, Didham RK, Fahrig L, Batáry P, Bengtsson J, Clough Y, Crist TO, Dormann CF, et al. 2012. Landscape moderation of biodiversity patterns and processes ‐ eight hypotheses. Biological Reviews 87: 661–685.

Uroy L, Ernoult A, Mony C. 2019. Effect of landscape connectivity on plant communities: a review of response patterns. Landscape Ecology 34: 203–225.

Vandenkoornhuyse P, Quaiser A, Duhamel M, Le Van A, Dufresne A. 2015. The importance of the microbiome of the plant holobiont. New Phytologist 206: 1196–1206.

Vannier N, Mony C, Bittebiere A-K, Michon-Coudouel S, Biget M, Vandenkoornhuyse P. 2018. A microorganisms’ journey between plant generations. Microbiome 6: 79.

Vannier N, Mony C, Bittebiere A-K, Theis KR, Rosenberg E, Vandenkoornhuyse P. 2019. Clonal Plants as Meta‐Holobionts. mSystems 4: e00213‐18.

Vašutová M, Mleczko P, López-García A, Maček I, Boros G, Ševčík J, Fujii S, Hackenberger D, Tuf IH, Hornung E, et al. 2019. Taxi drivers: the role of animals in transporting mycorrhizal fungi. Mycorrhiza 29: 413–434.

Voges MJEEE, Bai Y, Schulze-Lefert P, Sattely ES. 2019. Plant‐derived coumarins shape the composition of an Arabidopsis synthetic root microbiome. Proceedings of the National Academy of Sciences 116: 12558–12565.

Wagner MR, Lundberg DS, del Rio TG, Tringe SG, Dangl JL, Mitchell-Olds T. 2016. Host genotype and age shape the leaf and root microbiomes of a wild perennial plant. Nature Communications 7: 12151.

Wang J, Shi X, Zheng C, Suter H, Huang Z. 2021. Different responses of soil bacterial and fungal communities to nitrogen deposition in a subtropical forest. Science of The Total Environment 755: 142449.

Werner GDA, Kiers ET. 2015. Order of arrival structures arbuscular mycorrhizal colonization of plants. New Phytologist 205: 1515–1524.

Xu Q, Ling N, Quaiser A, Guo J, Ruan J, Guo S, Shen Q, Vandenkoornhuyse P. 2022. Rare Bacteria Assembly in Soils Is Mainly Driven by Deterministic Processes. Microbial Ecology 83: 137–150.

Xu Q, Vandenkoornhuyse P, Li L, Guo J, Zhu C, Guo S, Ling N, Shen Q. 2021. Microbial generalists and specialists differently contribute to the community diversity in farmland soils. Journal of Advanced Research: S2090123221002423.

Yuen J, Mila A. 2015. Landscape‐Scale Disease Risk Quantification and Prediction. Annual Review of Phytopathology 53: 471–484.

Zanne AE, Abarenkov K, Afkhami ME, Aguilar-Trigueros CA, Bates S, Bhatnagar JM, Busby PE, Christian N, Cornwell WK, Crowther TW, et al. 2020. Fungal functional ecology: bringing a trait‐based approach to plant‐associated fungi. Biological Reviews 95: 409–433.

Zhang J, Zhang B, Liu Y, Guo Y, Shi P, Wei G. 2018. Distinct large‐scale biogeographic patterns of fungal communities in bulk soil and soybean rhizosphere in China. Science of The Total Environment 644: 791–800.

Zuur AF, Ieno EN, Walker N, Saveliev AA, Smith GM. 2009. Mixed effects models and extensions in ecology with R. New York, NY: Springer New York.

