## Supplemental materials for "Ecological corridors homogenize plant root endospheric mycobiota"

**This PDF file includes:**

Supporting materials and methods

Supporting references

Supplementary Figures S1 to S13

Supplementary Tables S1 to S3

**Supporting materials and methods**

**Experimental design**

The mesocosms were filled with a homogeneous substrate of sand (20%) and agricultural silty clay soil (80%). To create the matrix, we planted 48 standardized clonal fragments of *B. pinnatum* (i.e., one ramet and one spacer) using a hexagonal planting pattern to guarantee balanced competitive interactions (Birch *et al.*, 2007). The matrix was created in June 2009, eight years before the *T. repens* patches were grown, to condition and homogenize the microbial reservoir. To restrict plant growth dynamics to the initially planted *B. pinnatum* individuals, we manually removed all mature flowers to prevent them from producing seeds. To create the patches, using the same batch of seeds, *T. repens* was germinated on sterile vermiculite in greenhouse conditions. During the indoor culture stage, the seedlings were watered with sterile water every second day and with a sterile hydroponic Hoagland solution (Arnon & Hoagland, 1940) once a week. After four weeks, in June 2017, the plantlets were transferred to the 20 mesocosms and formed the patches. Twenty plantlets were transplanted in a hexagonal pattern in each of the patches, and 10 plantlets were transplanted to the corridors with equal spacing between each plantlet. The patches were delimited by PVC circles, 15-cm tall and 40 cm in diameter. The corridors were 90 cm long and 15 cm wide and delimited by PVC edges 15-cm tall (Fig.1). Obstacles were placed on the surface of the ground to limit clonal growth of *T. repens* through runners and to prevent it from colonizing the *B. pinnatum* matrix. However, below the ground, roots of the two plant species could come into contact as no physical obstacles were placed under the soil surface.

Both *T. repens* and *B. pinnatum* plants in the mesocosms were watered twice a week in the dry seasons. Matrix vegetation was mowed twice a year, in spring and summer, to limit competition for light within *B. pinnatum* and *T. repens and* between the two species. We regularly manually removed *B. pinnatum* plantlets that were able to reach the *T. repens* patches by developing rhizomes and all the weeds growing in the mesocosms.

**Root sampling, DNA extraction and amplicon sequencing**

The root systems were carefully cleaned with tap water. For each sample, we then selected 100 mg of roots (fresh weight) from a representative subsample of the root system. The surface of the root was cleaned in a 2‰ Triton X100 solution for 5 mins, then rinsed twice with tap water, finally the roots were rinsed with sterile distilled water. The roots were then cut into small pieces and ground into powder. The total DNA of root associated microbiome was extracted from the root powder using a standard protocol with magnetic beads (Sbeadex mini plant kit, LGC Genomics) and an automated protocol (oKtopure robot platform, LGC Genomics) on the Gentyane platform(<https://www6.clermont.inrae.fr/umr1095_eng/Organisation/Experimental-Infrastructure/High-throughput-sequencing-genotyping>). DNA concentrations were normalized to 12.5 ng/μL after a fluorometric Hoechst DNA assay in preparation for subsequent next-generation sequencing. A specific 550 bp of the fungal small subunit (SSU) rRNA gene fragment, including the V4 and V5 regions, was amplified from 50 ng of extracted total DNA of root-associated microbiome using the primers SSU0817 (5’-TTAGCATGGAATAATRRAATAGGA-3’) and NS22B (5’-AATTAAGCAGACAAATCACT-3’) (Borneman & Hartin, 2000; Lê Van *et al.*, 2017). PCRs, performed with Illustra PuReTaq Ready-to-go (GE Healthcare, Chicago, IL, USA), contained 0.2 μM of primers in a final volume of 25 μL. The cycling regime was identical to the methods used by (Lê Van *et al.*, 2017). On 5’, the two primers contained the Illumina tails required a second round PCR and the amplicon identification necessary for multiplexing. Each amplicon was then purified (Agencourt AMPure) using a Bravo Automated Liquid Handling Platform (Agilent, Santa Clara, CA, USA), then quantified (Quant-iT PicoGreen dsDNA Assay Kit) and normalized to the same concentrations before pair-end sequencing (Miseq; Illumina, San Diego, CA, USA). Sequence data were analyzed using the FROGS pipeline (Escudié *et al.*, 2018). Data for 821 samples with 266 sequence clusters were organized in a contingency table (39 samples were discarded due to their low DNA concentration, low PCR product quality and quantity, or insufficient sequence depth) and the abundance of sequences per sample was rarefied by the lowest sequence depth (4 292) (Fig. S1).

**Supporting references**

**Arnon DI, Hoagland DR**. **1940**. Crop production in artificial culture solutions and in soils with special reference to factors influencing yields and absorption of inorganic nutrients. **50:463-485**.

**Birch CPD, Oom SP, Beecham JA**. **2007**. Rectangular and hexagonal grids used for observation, experiment and simulation in ecology. *Ecological Modelling* **206**: 347–359.

**Borneman J, Hartin RJ**. **2000**. PCR Primers That Amplify Fungal rRNA Genes from Environmental Samples. *Applied and Environmental Microbiology* **66**: 4356–4360.

**Escudié F, Auer L, Bernard M, Mariadassou M, Cauquil L, Vidal K, Maman S, Hernandez-Raquet G, Combes S, Pascal G**. **2018**. FROGS: Find, Rapidly, OTUs with Galaxy Solution (B Berger, Ed.). *Bioinformatics* **34**: 1287–1294.

**Lê Van A, Quaiser A, Duhamel M, Michon-Coudouel S, Dufresne A, Vandenkoornhuyse P**. **2017**. Ecophylogeny of the endospheric root fungal microbiome of co-occurring *Agrostis stolonifera*. *PeerJ* **5**: e3454.

**Supplementary figures**

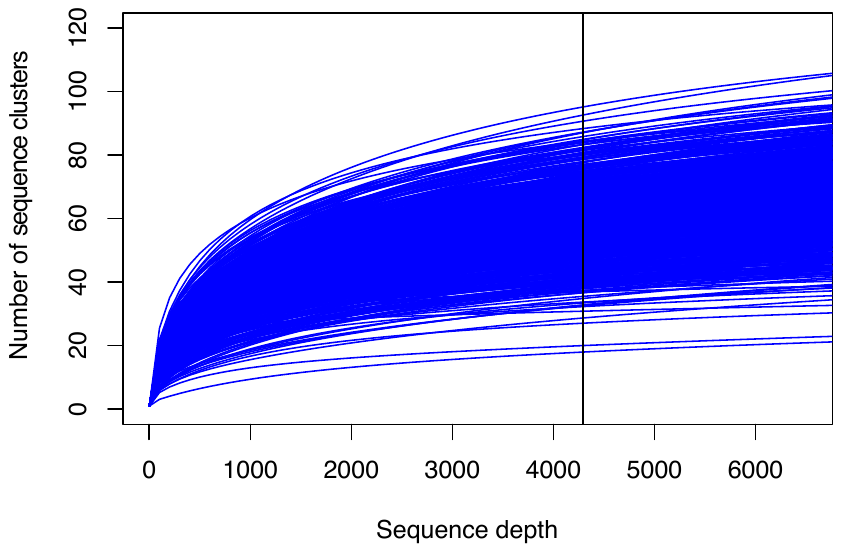

**Fig. S1.** **Rarefaction curves for sequence data of all samples.** Curves show the total number of sequence clusters detected relative to the sample size in number of sequences. The vertical line indicates the sample size used to rarefy the sequence data of all samples.

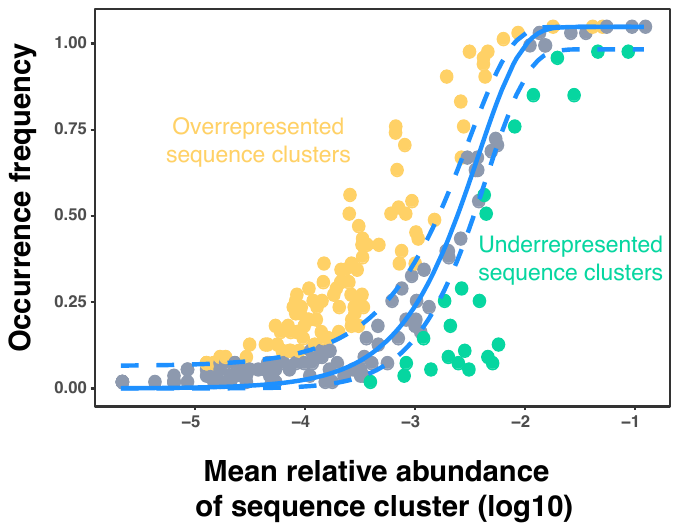

**Fig. S2. An example of Sloan’s neutral model:** the blue curve is the neutral model, the dashed blue curves are the 95% confidence intervals, the empty symbols inside the blue curves are neutral sequence clusters, which are stochastic, while the circles above and below the blue curves are overrepresented or underrepresented sequence clusters, respectively, which are deterministic.

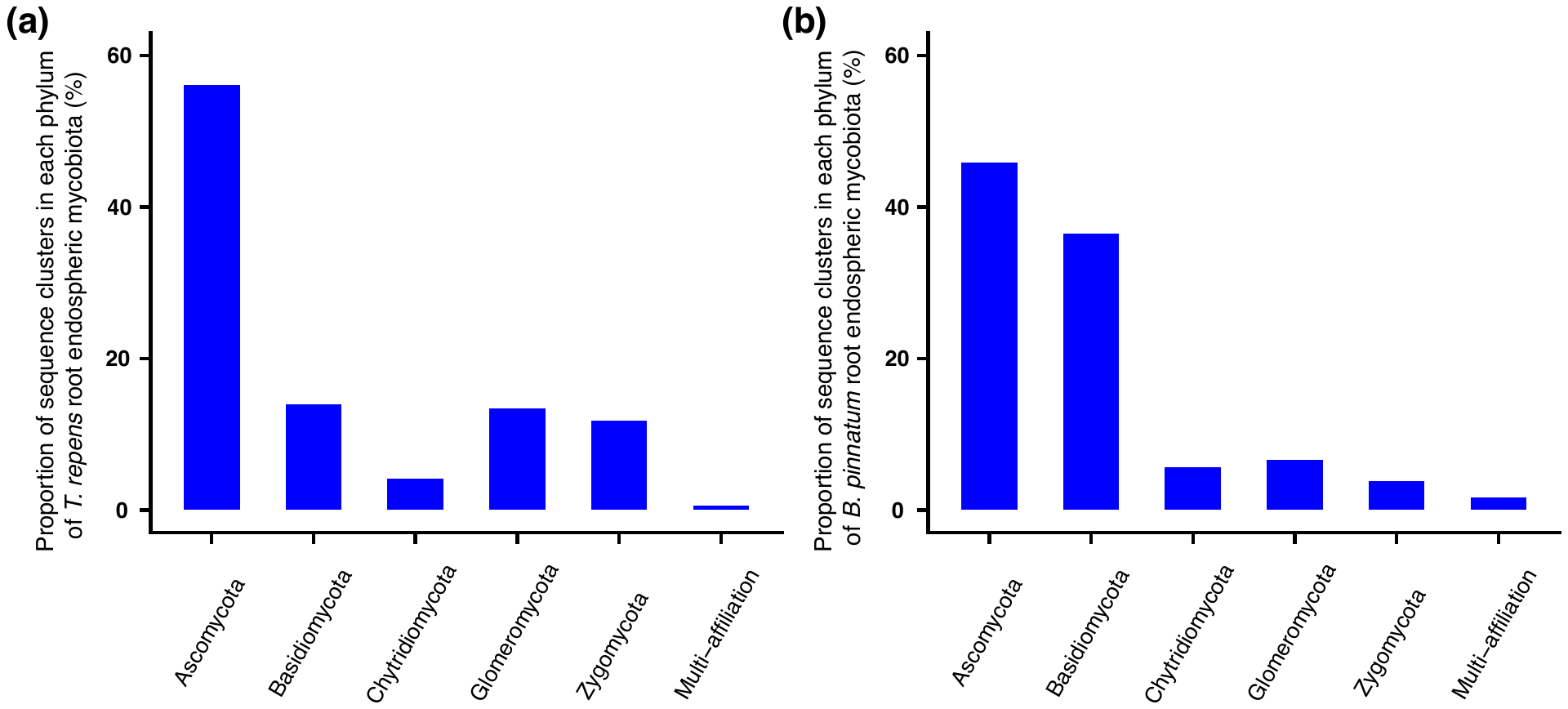

**Fig. S3.** **Proportion of each phylum in root endospheric mycobiota sequence clusters of *T. repens* and *B. pinnatum*.** (a) proportion of sequence clusters within each phylum (Ascomycota, Basidiomycota, Chytridiomycota, Glomeromycota and Zygomycota) of *T. Repens* root mycobiota; (b) proportion of sequence clusters within each phylum (Ascomycota, Basidiomycota, Chytridiomycota, Glomeromycota and Zygomycota) of *B. Pinnatum* root mycobiota.

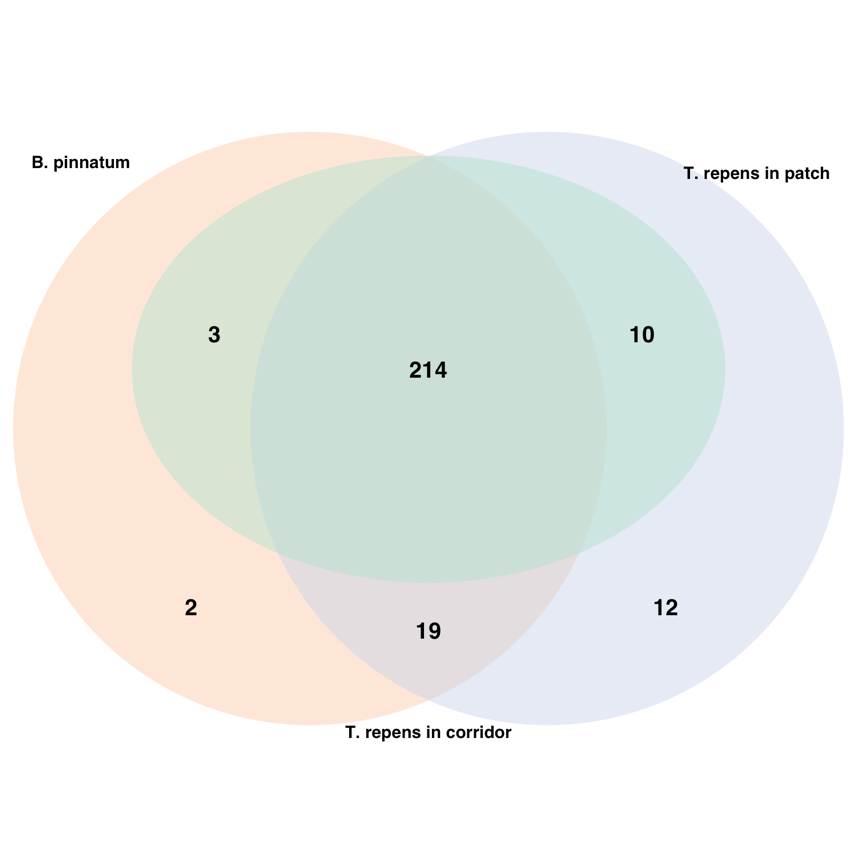

**Fig. S4. Venn diagram of shared and unique root endospheric mycobiota of *T. repens* in patches or corridors and *B. Pinnatum* across all sampling campaigns at individual scale.**

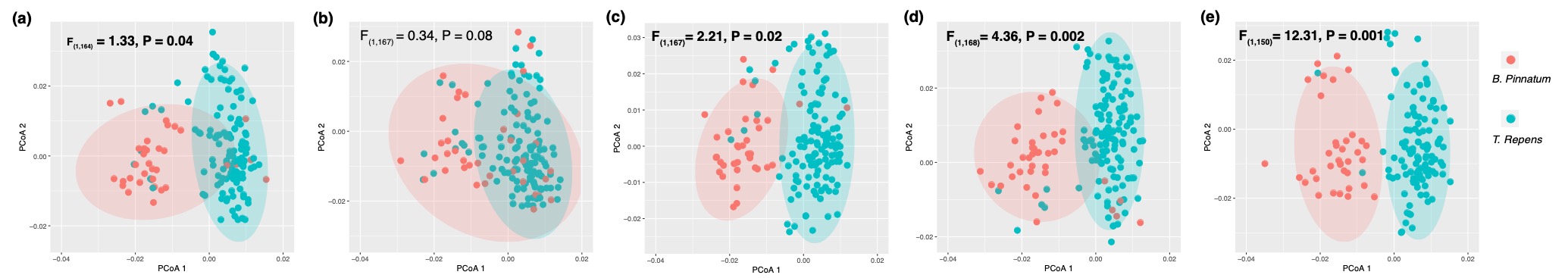

**Fig. S5. Root endospheric mycobiota community structure of *T. repens* and *B. Pinnatum* across all sampling campaigns at individual scale.** (a) plants sampled in October 2017; (b) plants sampled in May 2018; (c) plants sampled in June 2018; (d) plants sampled in October 2018; (e) plants sampled in May 2019. Ellipses in each figure show the 95% confidence interval. Statistics are indicated by F and P values, Significant results (P < 0.05) are in bold.

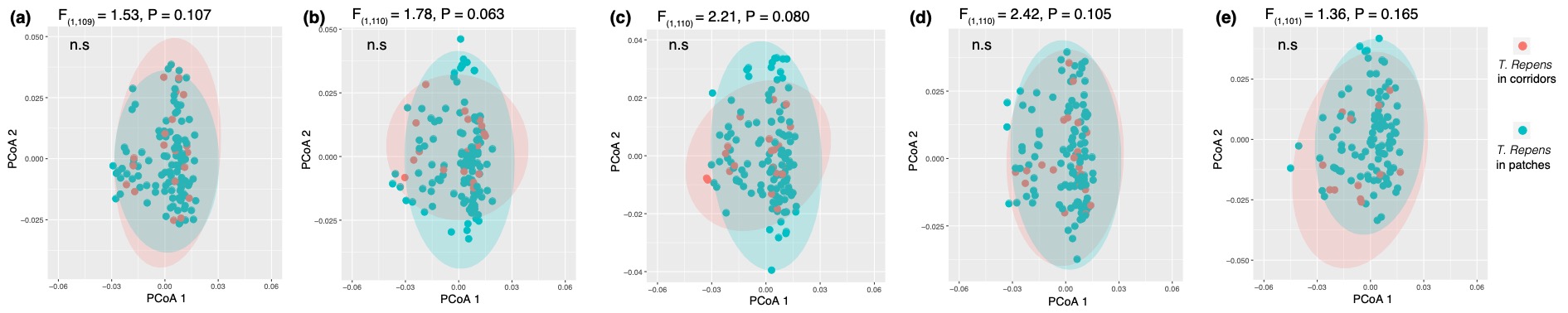

**Fig. S6. Root endospheric mycobiota community structure of *T. repens* from corridors and patches at individual scale.** (a) *T. repens* sampled in October 2017; (b) *T. repens* sampled in May 2018; (c) *T. repens* sampled in June 2018; (d) *T. repens* sampled in October 2018; (e) *T. repens* sampled in May 2019. Ellipses in each figure show the 95% confidence interval. Statistics are indicated by F and P values; n.s.: not significant

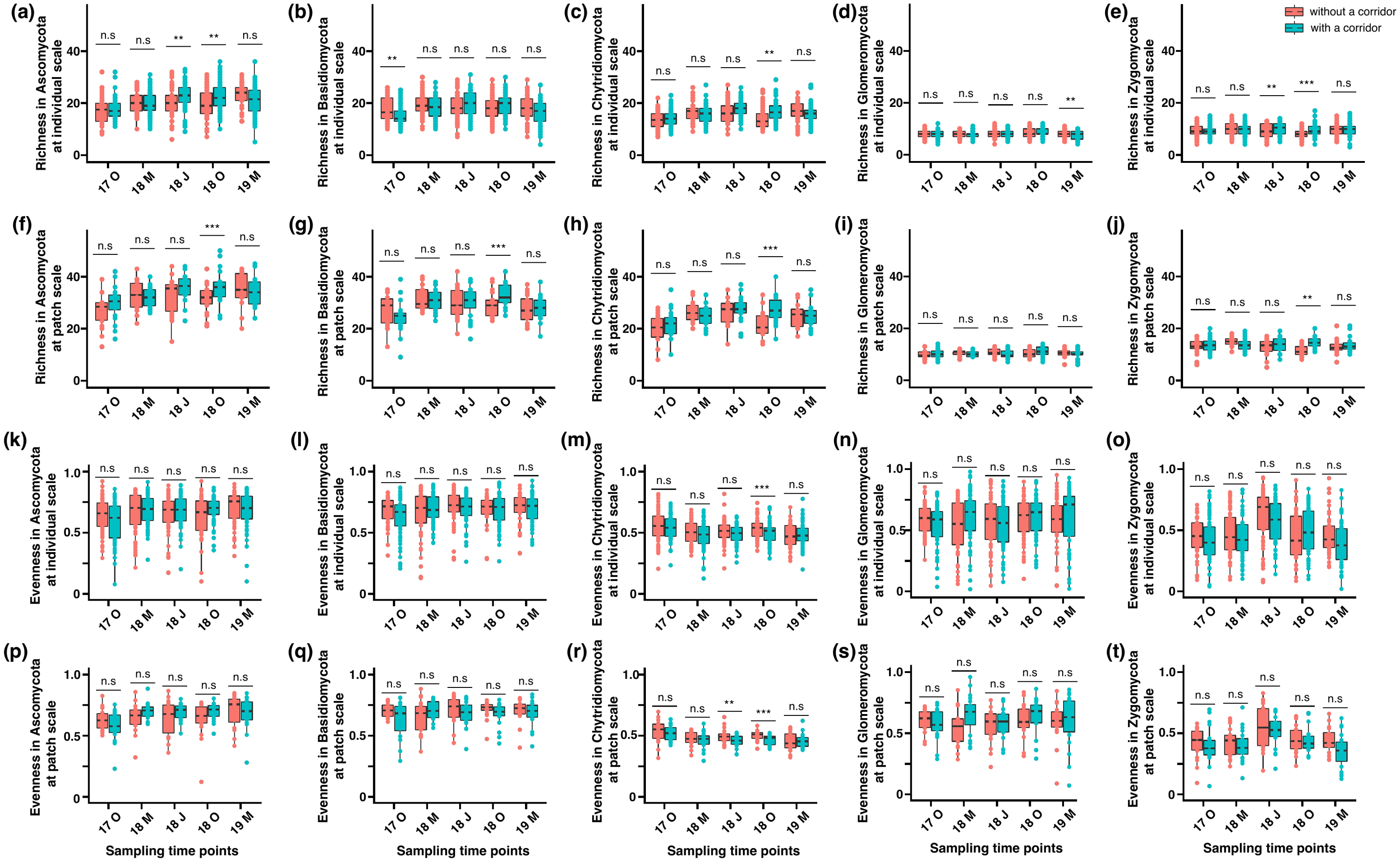

**Fig. S7.** **Root endospheric mycobiota diversity for each fungal phylum associated with *T. repens* in treatments with and without a corridor over five sampling campaigns at individual and patch scales.** Root endospheric mycobiota diversity was calculated as sequence cluster richness and Pielou’s evenness. Indices were calculated and tested at two different biological scales: at plant individual (i.e. one root sample; a-e; k-o) and patch (i.e. group of three *T. repens* root samples from the same patch pooled; f-j; p-t). (a) and (f) sequence cluster richness of root endospheric mycobiota in phylum Ascomycota at individual and patch scales, respectively; (b) and (g) sequence cluster richness of root endospheric mycobiota in phylum Basidiomycota at individual and patch scales, respectively; (c) and (h) sequence cluster richness of root endospheric mycobiota in phylum Chytridiomycota at individual and patch scales, respectively; (d) and (i) sequence cluster richness of root §endospheric mycobiota in phylum Glomeromycota at individual and patch scales, respectively; (e) and (j) sequence cluster richness of root endospheric mycobiota in phylum Zygomycota at individual and patch scales, respectively. (k) and (p) sequence cluster Pielou’s evenness of root endospheric mycobiota in phylum Ascomycota at individual and patch scales, respectively; (l) and (q) sequence cluster Pielou’s evenness of root endospheric mycobiota in phylum Basidiomycota at individual and patch scales, respectively; (m) and (r) sequence cluster Pielou’s evenness of root endospheric mycobiota in phylum Chytridiomycota at individual and patch scales, respectively; (n) and (s) sequence cluster Pielou’s evenness of root endospheric mycobiota in phylum Glomeromycota at individual and patch scales, respectively; (o) and (t) sequence cluster Pielou’s evenness of root endospheric mycobiota in phylum Zygomycota at individual and patch scales, respectively. 17 O: October 2017; 18 M: May 2018; 18 J: June 2018; 18 O: October 2018; 19 M: May 2019. Asterisks indicate the level of significance of the presence of a corridor on root sequence cluster richness: * 0.01 < P < 0.05; ** 0.001 < P < 0.01; *** P < 0.001; n.s.: not significant

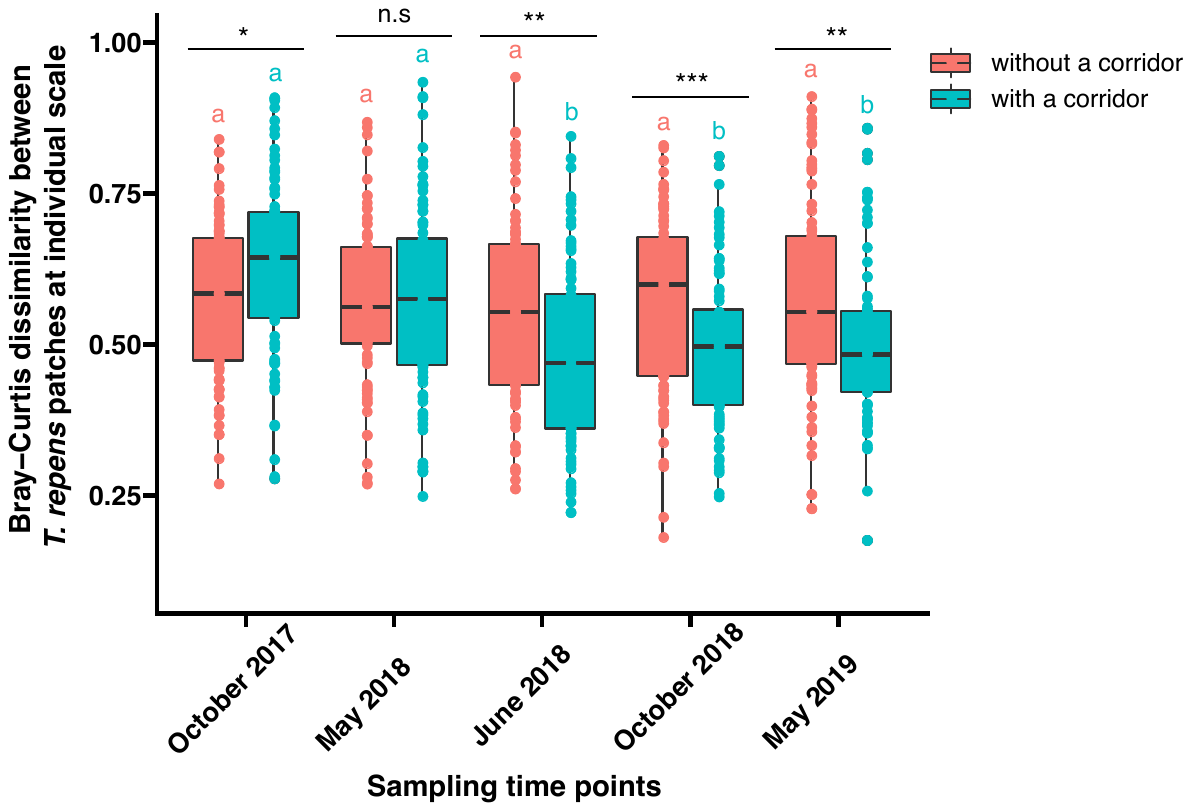

**Fig. S8.** **Effect of the presence of a corridor on *T. repens* root endospheric mycobiota dissimilarity at individual scale.** *T. repens* root endospheric mycobiota dissimilarity was calculated as Bray-Curtis dissimilarity between root mycobiota of pairwise *T. repens* patches from the same mesocosm with and without a corridor over five sampling campaigns at the individual scale. Nine possible pairs of Bray-Curtis dissimilarity between root mycobiota of three plant individuals in patch 1 and three plant individuals in patch 2 of the same mesocosm were calculated. Corridor effects were conducted via mixed linear models with the mesocosm as a random effect on Bray-Curtis dissimilarity, and the significance is shown by asterisks at the top of each bar: * 0.01 < P < 0.05; ** P < 0.01; *** P < 0.001; n.s.: not significant. Time effects were calculated via mixed linear models with the mesocosm as a random effect on Bray-Curtis dissimilarity along with sampling time points with and without a corridor, and the significance is shown by lowercase letters.

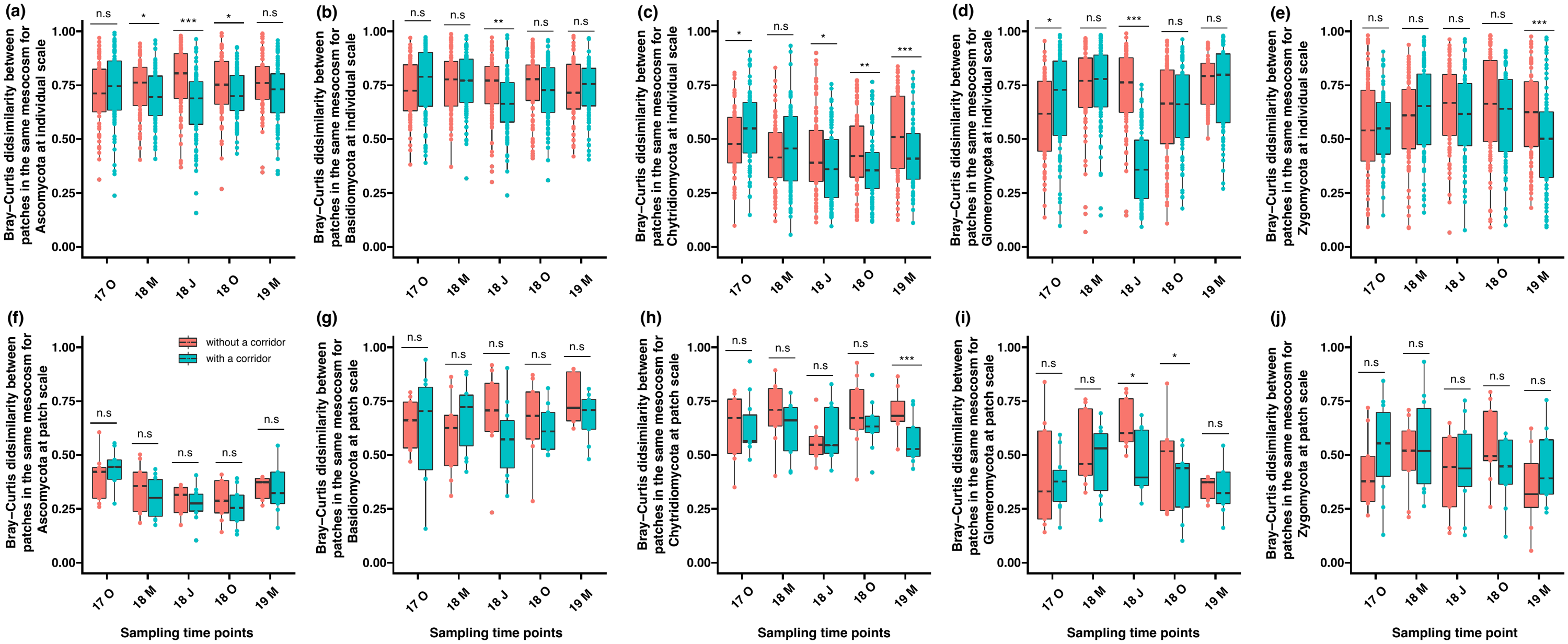

**Fig. S9.** **Effect of the presence of a corridor on *T. repens* root endospheric mycobiota dissimilarity of each phylum at individual and patch scales.** Indices were calculated and tested at two different biological scales: at plant individual (i.e. one root sample; a-e) and patch (i.e. group of three *T. repens* root samples from the same patch pooled; f-j). Dissimilarity was calculated as Bray-Curtis dissimilarity between root mycobiota for each phylum of pairwise *T. repens* patches from the same mesocosms with and without a corridor over five sampling campaigns at the individual and patch scale. (a) and (f) Bray-Curtis dissimilarity between root mycobiota for phylum Ascomycota at individual and patch scales, respectively; (b) and (g) Bray-Curtis dissimilarity between root mycobiota for phylum Basidiomycota at individual and patch scales, respectively; (c) and (h) Bray-Curtis dissimilarity between root mycobiota for phylum Chytridiomycota at individual and patch scales, respectively; (d) and (i) Bray-Curtis dissimilarity between root mycobiota for phylum Glomeromycota at individual and patch scales, respectively; (e) and (j) Bray-Curtis dissimilarity between root mycobiota for phylum Zygomycota at individual and patch scales, respectively. 17 O: October2017; 18 M: May 2018; 18 J: June 2018; 18 O: October 2018; 19 M: May 2019. Individual scale corridor effects were calculated via mixed linear models with the mesocosm as a random effect on Bray-Curtis dissimilarity, while patch scale corridor effects were calculated via t-tests with no random effects. Significant results have asterisks at the top of each bar: * 0.01 < P < 0.05; ** 0.001 < P < 0.01; *** P < 0.001; n.s./ not significant.

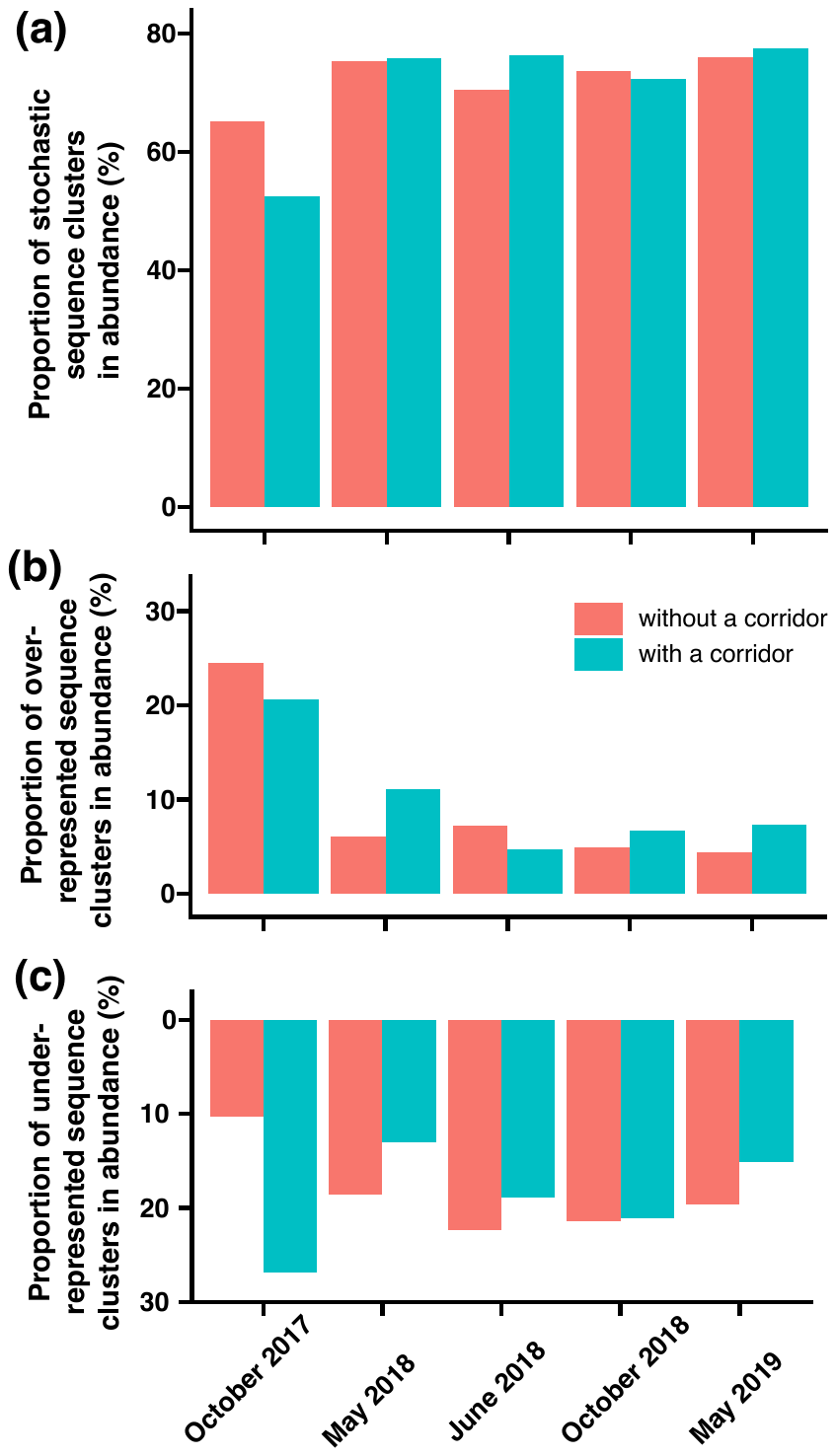

**Fig. S10. Summary of root endospheric mycobiota assembly patterns based on their abundance with and without corridors at five sampling time points.** (a) proportion of neutral sequence clusters among all the sequence clusters based on their abundance in both treatments, i.e. with and without a corridor; (b) proportion of overrepresented sequence clusters among all the sequence clusters based on their abundance in both treatments, i.e. with and without a corridor; (c) proportion of underrepresented sequence clusters among all the sequence clusters based on their abundance in both treatments, i.e. with and without a corridor.

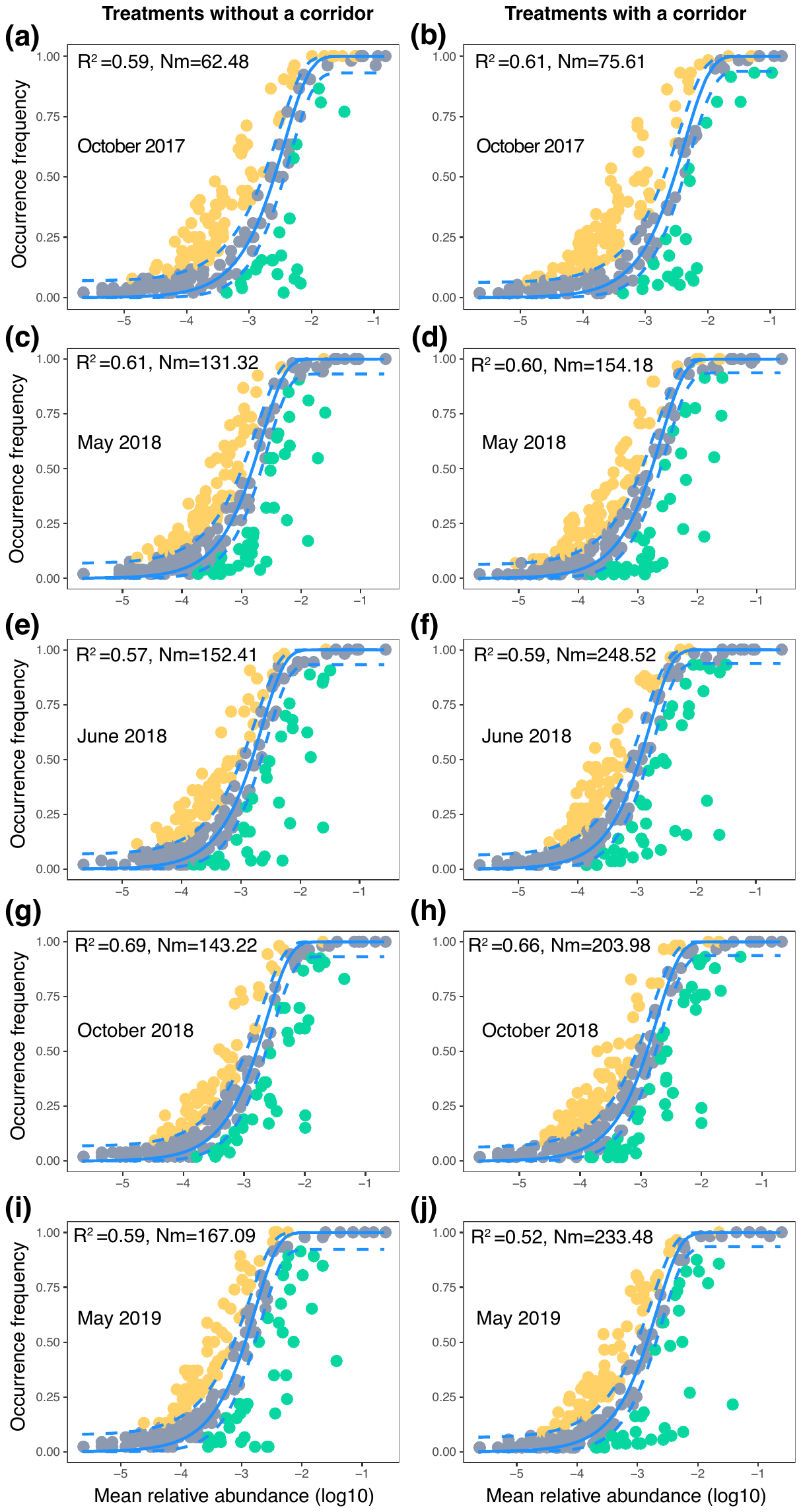

**Fig. S11. Sloan neutral community model for root endospheric mycobiota in treatments without (left panels) and with (right panels) corridors at each sampling time point.** (a) and (b) *T. repens* sampled in October 2017; (c) and (d) *T. repens* sampled in May 2018; (e) and (f) *T. repens* sampled in June 2018; (g) and (h) *T. repens* sampled in October 2018; (i) and (j) *T. repens* sampled in May 2019. The blue dashed curves in each panel represent the 95% confidence interval of the neutral model. The gray dots inside the dashed blue curves are neutral sequence clusters which are stochastic, while the dots above and below the blue curves are overrepresented or underrepresented sequence clusters, respectively, which are deterministic. Statistics of each model are shown as R^2^ and Nm in each panel: Rsqr represents the overall fit to each neutral model; N represents the metacommunity size, m represents the immigration rate, Nm is an estimate of the dispersal of root mycobiota communities between patches. More detailed statistics are included in Table S3.

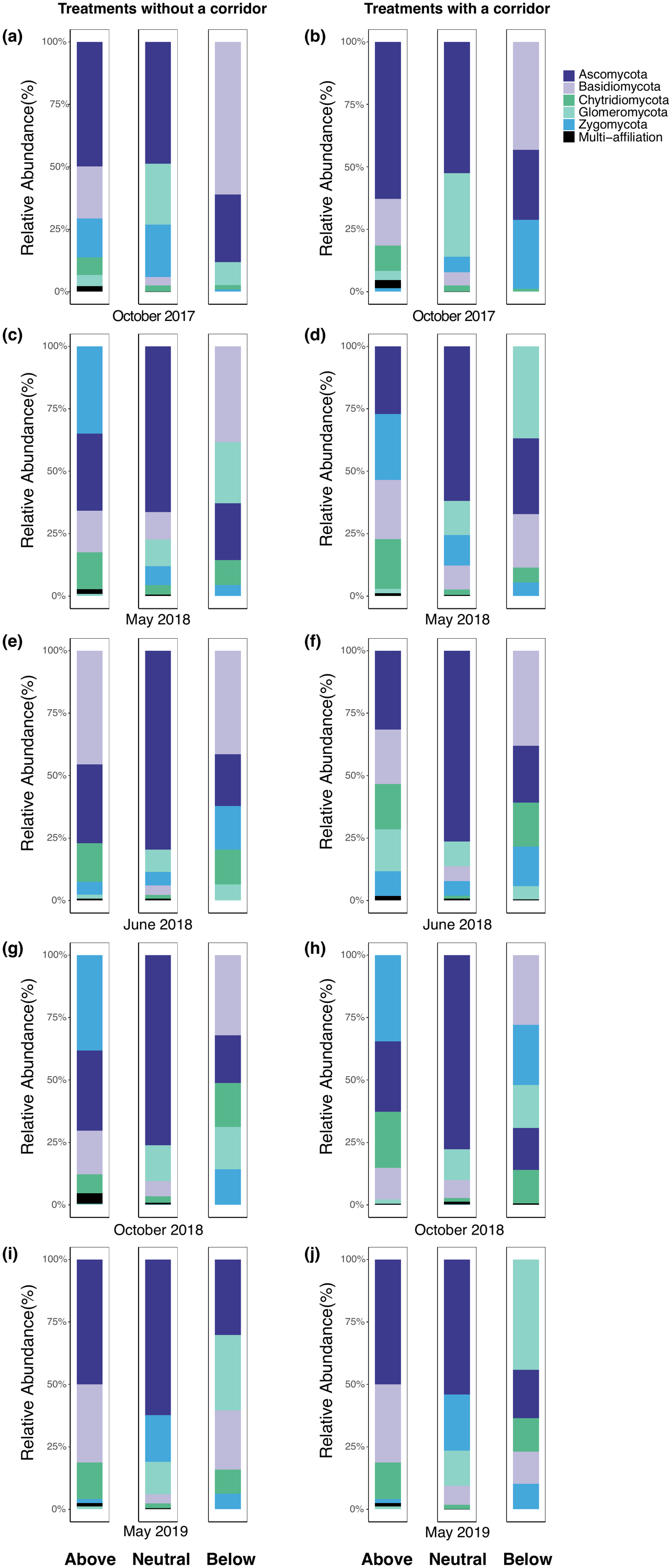

**Fig. S12. Taxonomic identity of the Sloan neutral community model for root endospheric mycobiota in treatments without a corridor (left panels) and with a corridor (right panels) at each sampling time point.** (a) and (b) *T. repens* sampled in October 2017; (c) and d) *T. repens* sampled in May 2018; (e) and (f) *T. repens* sampled in June 2018; (g) and (h) *T. repens* sampled in October 2018; (i) and (j) *T. repens* sampled in May 2019. In each panel, the left bar shows the taxonomic composition of overrepresented sequence clusters; the middle bar shows the taxonomic composition of neutral sequence clusters; the right bar shows the taxonomic composition of underrepresented sequence clusters.

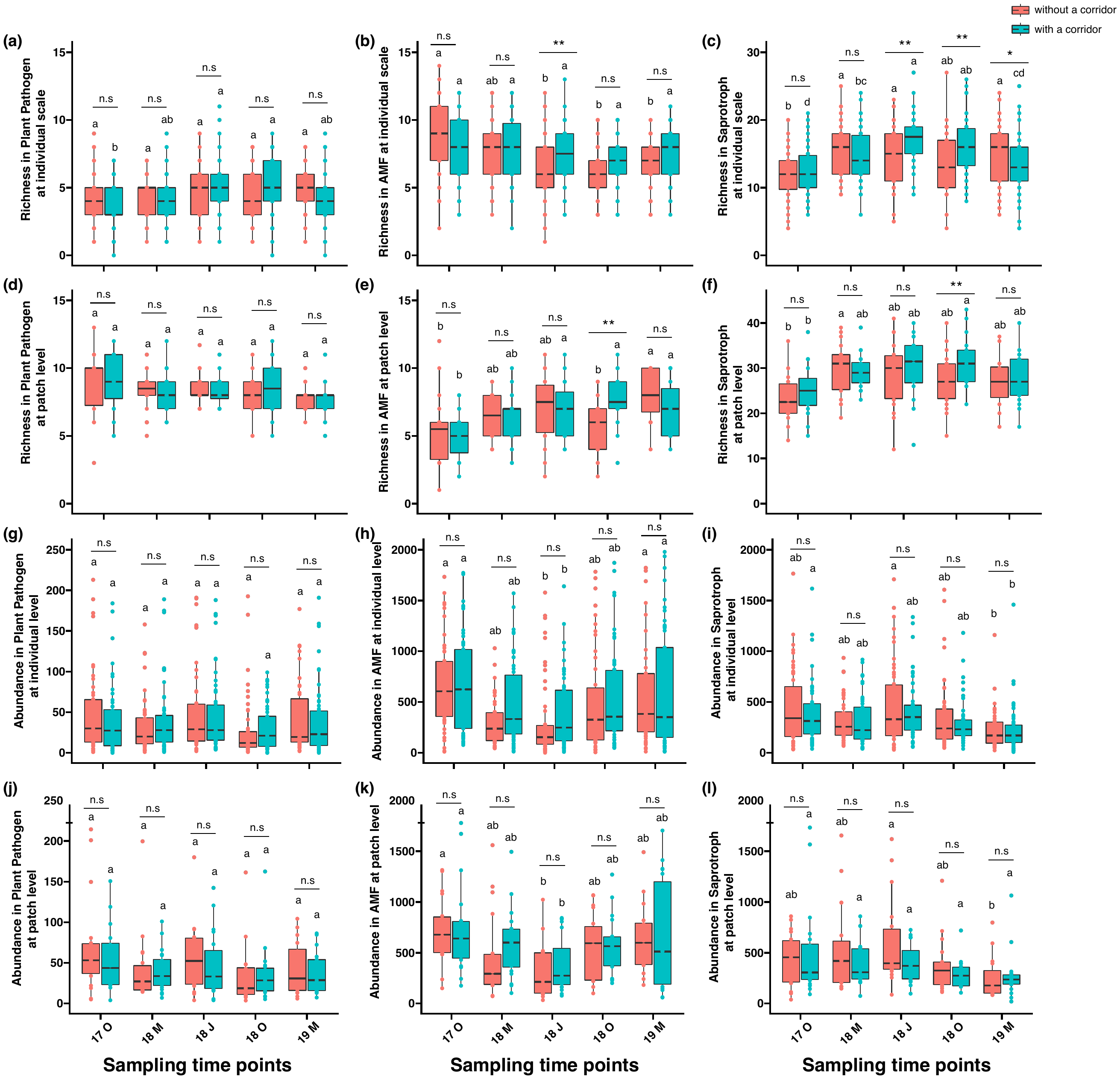

**Fig. S13. Dynamics of root endospheric mycobiota sequence cluster richness and abundance within each fungal guild over time in the treatments with a corridor and without a corridor at individual and patch scales.** (a) and (g) sequence cluster richness and abundance within a plant pathogen guild at individual scale, respectively; (b) and (h) sequence cluster richness and abundance within an arbuscular mycorrhizae (AMF) at individual scale, respectively; (c) and (i) sequence cluster richness and abundance within a Saprotroph guild at individual scale, respectively; (d) and (j) sequence cluster richness and abundance within a plant pathogen guild at patch scale, respectively; (e) and (k) sequence cluster richness and abundance within an arbuscular mycorrhizae (AMF) guild at patch scale, respectively; (f) and (l) sequence cluster richness and abundance within a Saprotroph guild at patch scale, respectively. On the x-axis, 17 O: October 2017; 18 M: May 2018; 18 J: June 2018; 18 O: October 2018; 19 M: May 2019. Pairwise comparisons were conducted. Asterisks at the top of each bar: * 0.01 < P < 0.05; ** P < 0.01; n.s.: not significant. Multiple group comparisons were conducted across sampling campaigns with and without corridors, and are indicated by lowercase letters.

**Supplementary tables**

**Table S1.** **Effects of the presence of a corridor, time, and their interaction on root endospheric mycobiota composition of *Trifolium repens* at individual and patch scales.** The effects were tested with a PERMANOVA analysis with the adonis function in R. Significant results (P < 0.05) are shown in bold.

| Different scales /Parameters | Root mycobiota composition at individual scale | | | Root mycobiota composition at patch scale | | |
| --- | --- | --- | --- | --- | --- | --- |
|  | df | F. Model | P | df | F. Model | P |
| Presence of a corridor | 1 | 2.90 | **0.001** | 1 | 1.43 | 0.070 |
| Sampling time point | 4 | 12.82 | **0.001** | 4 | 1.34 | **0.015** |
| Interactive effect | 4 | 1.41 | **0.004** | 4 | 1.57 | **0.001** |
| Residual | 535 |  |  | 175 |  |  |

**Table S2. Statistics for 10 Sloan Neutral Models.** m denotes the immigration rate; m.ci the fitting model parameter m using non-linear least squares; R^2^ represents the overall fit to the neutral model; RMSE: Root Mean Squared Error; AIC: Akaike’s Information Criterion; BIC: Bayesian information criterion; N stands for the metacommunity size; Samples for the number of samples; Richness for sequence cluster richness; Detect for the limit of detection; Nm is an estimate of dispersal of root mycobiota communities between patches with or without the presence of a corridor.

| **Corridor** | **Time** | **m** | **m.ci** | **R^2^** | **RMSE** | **AIC** | **BIC** | **N** | **Samples** | **Richness** | **Detect** | **Nm** |
| --- | --- | --- | --- | --- | --- | --- | --- | --- | --- | --- | --- | --- |
| without | October 2017 | 0.01 | 0.00 | 0.59 | 0.19 | -110.21 | -103.36 | 4290 | 58 | 227 | 0.0002 | 62.48 |
| with | October 2017 | 0.02 | 0.00 | 0.61 | 0.19 | -103.47 | -96.67 | 4290 | 52 | 221 | 0.0002 | 75.61 |
| without | May 2018 | 0.03 | 0.01 | 0.61 | 0.18 | -132.72 | -125.71 | 4290 | 58 | 245 | 0.0002 | 130.32 |
| with | May 2018 | 0.04 | 0.01 | 0.60 | 0.19 | -122.15 | -115.11 | 4288 | 53 | 250 | 0.0002 | 154.18 |
| without | June 2018 | 0.04 | 0.01 | 0.57 | 0.19 | -107.43 | -100.49 | 4290 | 53 | 237 | 0.0002 | 152.41 |
| with | June 2018 | 0.06 | 0.01 | 0.59 | 0.20 | -96.31 | -89.36 | 4290 | 58 | 238 | 0.0002 | 248.52 |
| without | October 2018 | 0.03 | 0.01 | 0.69 | 0.17 | -168.61 | -161.71 | 4195 | 53 | 233 | 0.0002 | 143.22 |
| with | October 2018 | 0.05 | 0.01 | 0.66 | 0.17 | -152.36 | -145.41 | 4288 | 58 | 238 | 0.0002 | 203.98 |
| without | May 2019 | 0.04 | 0.01 | 0.59 | 0.19 | -109.37 | -102.43 | 4234 | 56 | 237 | 0.0002 | 167.09 |
| with | May 2019 | 0.05 | 0.01 | 0.52 | 0.21 | -65.64 | -58.77 | 4288 | 46 | 229 | 0.0002 | 233.48 |

**Table S3.** **Taxonomic identifications and ecological functions of 138 root endospheric sequence clusters.** Sequence clusters with a yellow background were assigned with functional information to the Fun^fun^ 0.0.3 database, while sequence clusters with a green background were assigned with functional information to the FungalTraits 1.2 database. The rest of sequence clusters were assigned with functional information to the FUNGuild database.
